# Non-clinical safety of GRAd vector-based COVID-19 and HIV vaccines supports a platform regulatory approach

**DOI:** 10.1101/2025.11.24.689034

**Authors:** Reji Paalangara, Stephanie Gohin, Alexis Menard, Charlotte Amy, Wahiba Berrabah, Alexandra Rogue, Matthew A Getz, Aljawharah Alrubayyi, Simone Battella, Angelo Raggioli, Michela Gentile, Anthea Di Rita, Alessia Noto, Giuseppina Miselli, Fabiana Grazioli, Federico Napolitano, Dhurata Sowcik, Marco Soriani, Benjamin Chmielewski, Lebohang Molife, Vincent Muturi-Kioi, Azure T Makadzange, Gaurav D Gaiha, Philippe Ancian, Jim Ackland, Antonella Folgori, Stefano Colloca, Stefania Capone

**Affiliations:** IAVI, New York, NY 10004, United States; Charles River Laboratories Evreux, 27930 Miserey, France; Ragon Institute of Mass General Brigham, MIT, and Harvard, Cambridge, MA 02139, USA; ReiThera srl, 00128 Rome, Italy; Mutala Trust, Vainona, Harare, Zimbabwe; Division of Gastroenterology, Massachusetts General Hospital and Harvard Medical School, Boston, MA 02115, USA; Program in Health Sciences and Technology, Harvard Medical School, Massachusetts Institute of Technology, Cambridge, MA 02115, USA; Global BioSolutions, Vermont, Victoria 3133, Australia

## Abstract

The rapid development of safe and efficacious vaccines is often hindered by extensive, mandated non-clinical safety evaluations in animals. Here we present the complete non-clinical studies for two investigational vaccines based on the GRAd platform, a gorilla-derived group C adenoviral vector. When administered intramuscularly, GRAd-COV2 and GRAdHIVNE1 were well tolerated. Studies in rats and rabbits showed localized distribution and transient, non-adverse inflammatory responses, while successfully inducing expected immune responses to their respective antigens. Notably, both vaccines demonstrated a consistent safety profile despite transgene and backbone differences, comparable to other replication-defective adenoviral vectors. The established non-clinical safety profile of the GRAd platform provides a robust foundation for a more efficient and streamlined regulatory pathway. By leveraging this prior knowledge, future GRAd-based vaccines can achieve accelerated clinical development while fully adhering to the ethical principles of replacement, reduction, and refinement of animal use in research.

## Introduction

Vaccines have provided significant protection from serious illnesses such as smallpox, polio, tetanus, diphtheria, pertussis, measles, and rubella for decades ^1^. More recently vaccines have prevented millions of deaths from COVID-19 ^2,3^. The primary mechanism of action of a vaccine is to deliver antigens to the human immune system triggering an immune response that can help prevent serious diseases. Traditional vaccine types include live attenuated or inactivated whole pathogens and purified or recombinant subunits, such as proteins, polysaccharide conjugates and toxoids, often requiring adjuvants for optimal efficacy. These approaches have proven successful to combat deadly pathogens, but with the downside of complex propagation, inactivation/killing and manufacturing processes often requiring years to be optimized and fully developed for each vaccine indication, limiting the opportunities for global disease prevention ^4^.

Modern vaccine platforms are based on standardized, reusable “modular” technologies where a carrier (i.e. nucleic acid, viral vector, or nanoparticle system) can be used interchangeably for different diseases by simply substituting the antigenic component ^5^. Such pathogen-agnostic platforms offer several advantages including reduced development cost and time due to well established manufacturing processes that can be quickly adapted for different indications. In addition, as novel vaccine candidates utilizing a given platform advance through development, the accumulating non-clinical and clinical safety data progressively refine our understanding of the platform’s safety and efficacy profile. Viral vectors and lipid nanoparticles have been used successfully to deliver genetic material (viral DNA or mRNA) encoding for a target vaccine antigen, allowing the host’s cellular machinery to produce the antigen in a physiologically relevant manner. This mechanism is especially crucial for eliciting robust MHC class I-restricted CD8^+^ T cell responses ^6^.

Of the initial effective COVID-19 vaccines approved within less than one year from the first identification of SARS-CoV-2 as the causative agent of the outbreak, six were based on established gene delivery vaccine platforms: either mRNA formulated in lipid nanoparticles (Moderna and Pfizer/BioNTech) or adenoviral vectors (Oxford/AstraZeneca, Janssen, Gamaleya and CanSino) ^7^. The COVID-19 pandemic, with all the associated major global health, societal and economic disruptions, has prompted the recognition of the need to accelerate the development of safe, effective, and equitable vaccines for preparedness against epidemic and pandemic threats. Similarly, for complex pathogens relevant for Global Health such as HIV, which is prone to extremely rapid mutation, the expedited assessment of innovative vaccine immunogens/combinations in small size experimental medicine clinical trials to enable subsequent iterations would be extremely helpful ^8^. However, the length of standard required non-clinical safety assessment to support regulatory approval of novel vaccine candidates for human clinical studies is not suited for such complex development scenarios, and the possibility to undertake a “platform approach” for non-clinical safety of well-studied vaccine platform technologies is an attractive perspective.

Replication-defective adenoviral (Ad) vectors, derived from human and non-human primates, are a well-characterized platform noted for potent immunogenicity, established safety, and manufacturing scalability^9^. Vaccines leveraging this mature platform have undergone extensive pre-clinical and clinical studies, with recent demonstrated efficacy leading to their approval and global deployment for diseases like Ebola and COVID-19. GRAd is a novel gorilla adenoviral species C strain isolated and successfully implemented as vector for a COVID-19 vaccine candidate. The GRAd-COV2 vaccine was proven safe and highly immunogenic in animal models ^10^ and in clinical trials up to phase 2 ^11–13^. The ability to induce a potent, broadly directed, polyfunctional and durable T cell response to the encoded SARS-CoV-2 Spike, especially of the CD8^+^ subtype, prompted the selection of GRAd as vector platform system for GRAdHIVNE1, an innovative T-cell inducing vaccine for HIV encoding highly networked, mutationally constrained CD8^+^ epitopes representing regions of vulnerability of the HIV virus ^14^. GRAdHIVNE1 has recently entered phase 1 clinical testing in people living without and with HIV in Southern African countries (NCT06617091). Here, we present the toxicity and biodistribution studies conducted with intramuscularly administered GRAd-COV2 and GRAdHIVNE1, in alignment with regulatory expectations for first-in-human vaccine trials. We show in detail their shared safety profiles despite the diverse nature of the encoded transgenes and subtle differences in the viral backbone. We also discuss our experimental findings in the context of published literature for other investigational or approved vaccines based on different replication-defective Ad vectors, highlighting the notable commonalities.

Finally, we present the scientific rationale supporting a limited non-clinical evaluation strategy for GRAd-based vaccines, according to a “platform approach”. This strategy is proposed to rapidly advance investigational vaccine candidates into early-phase clinical trials, substantially accelerating development without compromising safety. Concurrently, it reduces the use of laboratory animals in line with the 3R principles (Replacement, Reduction, and Refinement).

## Results

### Common and divergent features of GRAd-COV2 and GRAdHIVNE1 vaccines

The main commonalities and differences of GRAd-COV2 and GRAdHIVNE1 are summarized in Table 1. Both investigational vaccines are based on the gorilla adenovirus group C serotype, GRAd32, with a deletion of the adenoviral genomic E1 region to abolish replication capacity. The E3 region has also been deleted (ΔE3) to create sufficient space in the viral genome for insertion of large foreign antigens. The transgene expression cassette is inserted in the deleted E1 region, with identical regulatory elements, and leftward oriented transcription. Essentially, once produced and packaged, the GRAd-COV2 and GRAdHIVNE1 vaccine viral particles are expected to be identical in their capsid protein composition and resulting cell tropism/receptor usage. However, the two vaccines differ in the engineering of their E4 genome region. GRAd-COV2 has the E4 region deleted and replaced by Human Ad5 E4 orf6 (denominated “backbone c”), while for GRAdHIVNE1 the E4 region is chimeric, with autologous GRAd32 orf 1 to 3, and orfs 4-6 and 6/7 from Human Ad5 (“backbone b2”). The two backbones have different properties, and were purposely selected to increase productivity for the GRAd-COV2 vaccine, that was intended for global use in a pandemic setting, and increase thermostability at 4°C for GRAdHIVNE1 vaccine, for best storage and deployment in a low- and middle-income countries (LMIC) where disease prevention can be most impactful.

**Table 1:**
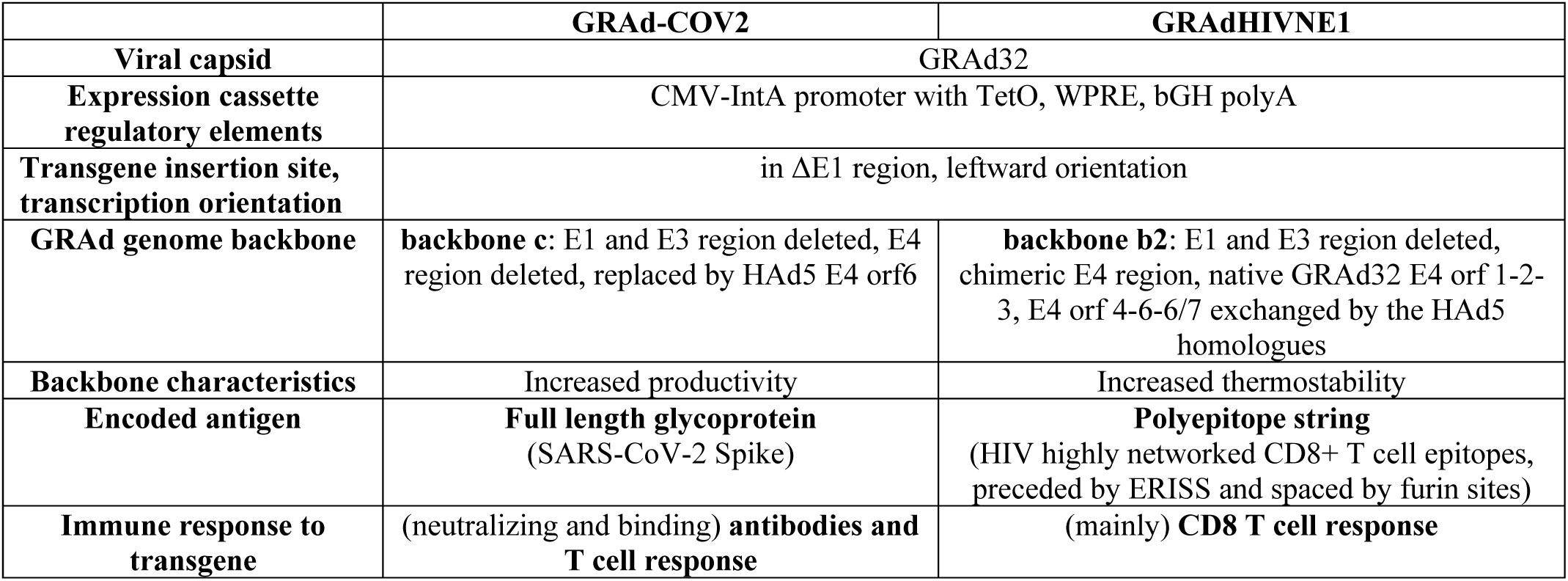
GRAd vaccines commonalities and differences.

The encoded antigens differ not only in their pathogens of origin (SARS-CoV-2 and HIV) but also in their nature and design: GRAd-COV2 encodes for a full length viral glycoprotein (Spike) stabilized in pre-fusion ^10^, while GRAdHIVNE1 encodes for a synthetic polyepitopic string assembled from HIV clades B and C regions enriched in highly networked CD8^+^ T cell epitopes, each spaced by furin cleavage sites for optimal processing and preceded by an endoplasmic reticulum targeting signal sequence (ERISS) signal sequence for endoplasmic reticulum (ER) translocation (manuscript in preparation). Therefore, the two transgenes follow different processing and antigen presentation pathways and are in fact designed to induce either neutralizing antibodies and T cell responses (GRAd-COV2) or mostly CD8^+^ T cell responses (GRAdHIVNE1).

### Localized biodistribution of GRAd vaccines in rats

The study design (single intramuscular administration in the quadriceps) and the rat as animal system were identical for the two vaccines, with a marginal difference in the administered dosage (2 × 10^10^ vp and 3.3 × 10^10^ vp) and injected volume (100µL or 200µL) for GRAd-COV2 and GRAdHIVNE1, respectively. Both vaccines were well tolerated; there were no mortalities, no clinical signs, no effects on body weight and food consumption, and no local reaction at injection site recorded during the *in vivo* phase of both studies. At necropsy, macroscopic findings identified included bilateral enlargement of draining inguinal and iliac lymph nodes with increased severity on the side of injection (right), decreasing over time. This was associated with increased inguinal (GRAd-COV2 only) and iliac lymph node weights, recorded in treated animals at all time-points. These findings are suggestive of immune activation upon vaccination.

GRAd-COV2 and GRAdHIVNE1 genomes were readily quantified in DNA extracted from the majority of injection sites and lymph node samples (Fig. 1 a-b and Supplementary tables 2 and 3), with the highest levels at day 2 and decreasing thereafter. Viral genomes were still detectable and quantifiable at 1 or 2 logs lower in both anatomical compartments at end of study (day 49), consistent with previous experience for replication defective Ad vectors.

**Figure 1.**
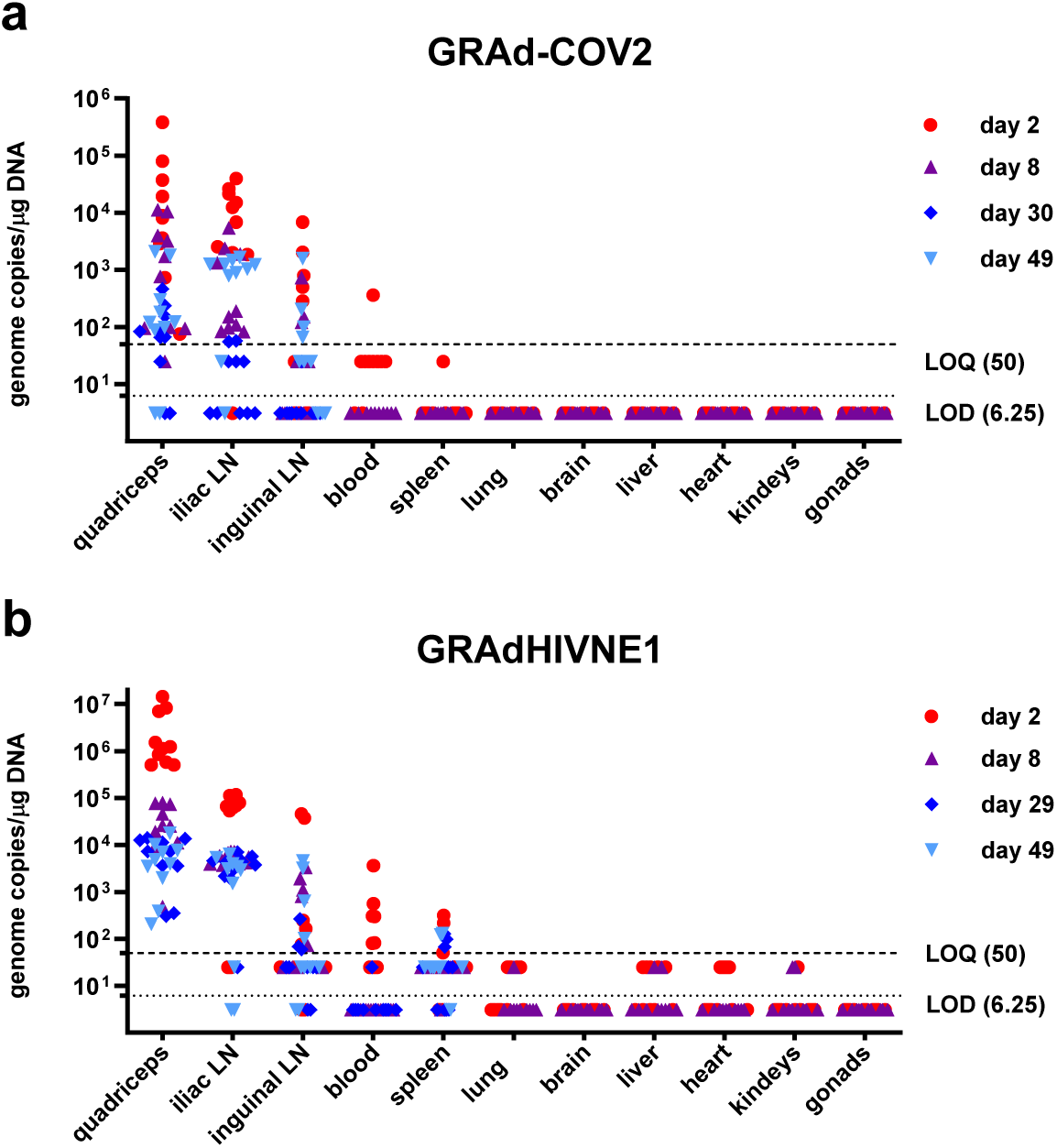
GRAd-COV2 and GRAdHIVNE1 biodistribution in rats. GRAd-COV2 (a) and GRAdHIVNE1 (b) genomes detected by qPCR in DNA extracted from indicated organs 2, 8, 29/30 and 49 days after a single intramuscular administration in the quadriceps, as highlighted by different symbols and colors. N=10 vaccinated animals at each time point. Data are expressed as genome copies per μg of DNA. Limit of detection (LOD) and lower limit of quantification (LOQ), established within assay validation, were set at 6.25 and 50 copies per 1μg of tested DNA respectively, and shown as dotted and dashed lines. Experimental samples resulting below LOD or LOQ were assigned values corresponding to one half of the extrapolated LOD and LOQ/μg of DNA (3.12 and 25 copies respectively). No GRAd-COV2 or GRAdHIVNE1 genomic DNA was quantified in any samples from control animals injected with saline, and thus are not reported in the graphs.

Both vaccines genomes were not quantified in any brain, heart, kidney, liver, lung, and gonad samples on days 2 and 8, and were not tested at later time points. Genome copies were quantified sporadically and at very low levels (361 copies/μg for GRAd-COV2 and up to 3490 copies/μg DNA for GRAdHIVNE1) in blood samples on day 2 only, more markedly for GRAdHIVNE1 (GRAd-COV2: 1 positive animal/10; GRAdHIVNE1: 7 positive/10); GRAdHIVNE1 only was quantified in spleen of few animals up to day 49, possibly indicative of translocation from the peripheral tissue and blood via the reticuloendothelial system.

### GRAd vaccines are well-tolerated in rabbit toxicity studies

In agreement with the Italian Regulatory agency (AIFA), a short duration single dose (SD-acute) toxicity study with limited read outs was conducted for GRAd-COV2 to enable rapid initiation of phase 1 clinical trial in the SARS-CoV-2 pandemic context, followed by a standard repeated dose (RD) toxicity and local tolerance study while the phase 1 trial was ongoing. For GRAdHIVNE1, a RD toxicity and local tolerance study was executed prior to the initiation of the clinical trial.

The common dose level of 1 × 10^11^ vp selected for all toxicity studies corresponds to one half of the highest human dose (2 × 10^11^ vp) intended for both vaccine candidates in clinical trials. Dosing 1 × 10^11^ vp in a ∼3 kg rabbit provided at least 10-fold Margin of Safety on a vp/Kg basis to the highest human dose, as appropriate.

For reference, Figures 2 and 3 show relevant in-life and clinical pathology parameters for the GRAd-COV2 SD toxicity study, since sampling and observation schedule was more extensive and informative in the week after the first administration. Similar findings were described for the RD toxicity studies of both GRAd-COV2 and GRAdHIVNE1 vaccines when sampling schedule allowed comparison, and any relevant discrepancies are discussed in the text. Supplementary tables 4 to 13 (for GRAd-COV2) and 14 to 19 (for GRAdHIVNE1) report parameters with significant changes in the RD toxicity studies.

**Figure 2.**
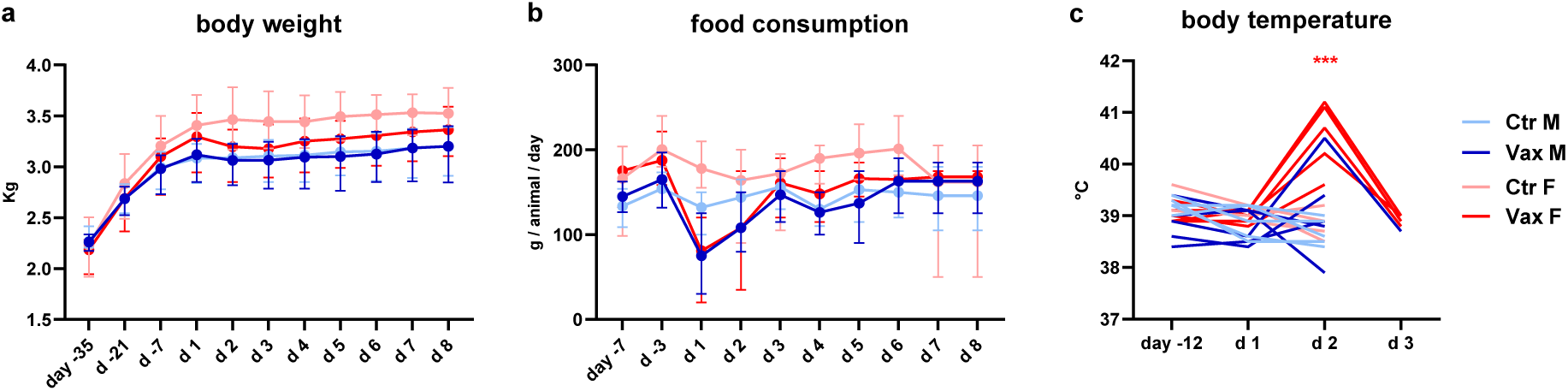
*in life* parameters in rabbits receiving a single saline or GRAd-COV2 administration. (a) absolute body weight, expressed in Kg and (b) food consumption, expressed in grams per animal per day, are reported for the duration of study follow up, before and up to 7 days after the GRAd-COV2 intramuscular administration (d1). Data represent the mean with range per study day of each treatment group. (c) rectal body temperature, expressed in degree Celsius (°C) measured in individual animals before (day -12), on day of GRAd-COV2 administration (d1) and until resolution. Throughout, the color code of the curves indicates treatment group assignment, either control male (light blue) and female (pink) animals, or GRAd-COV2 vaccinated male (blue) and female (red) animals. Statistically significant differences from controls of same sex, according to the statistical analysis decision tree, are indicated with blue (male) or red (female) asterisks: *(p=<0.05); **(p=<0.01); ***(p=<0.001)

**Figure 3.**
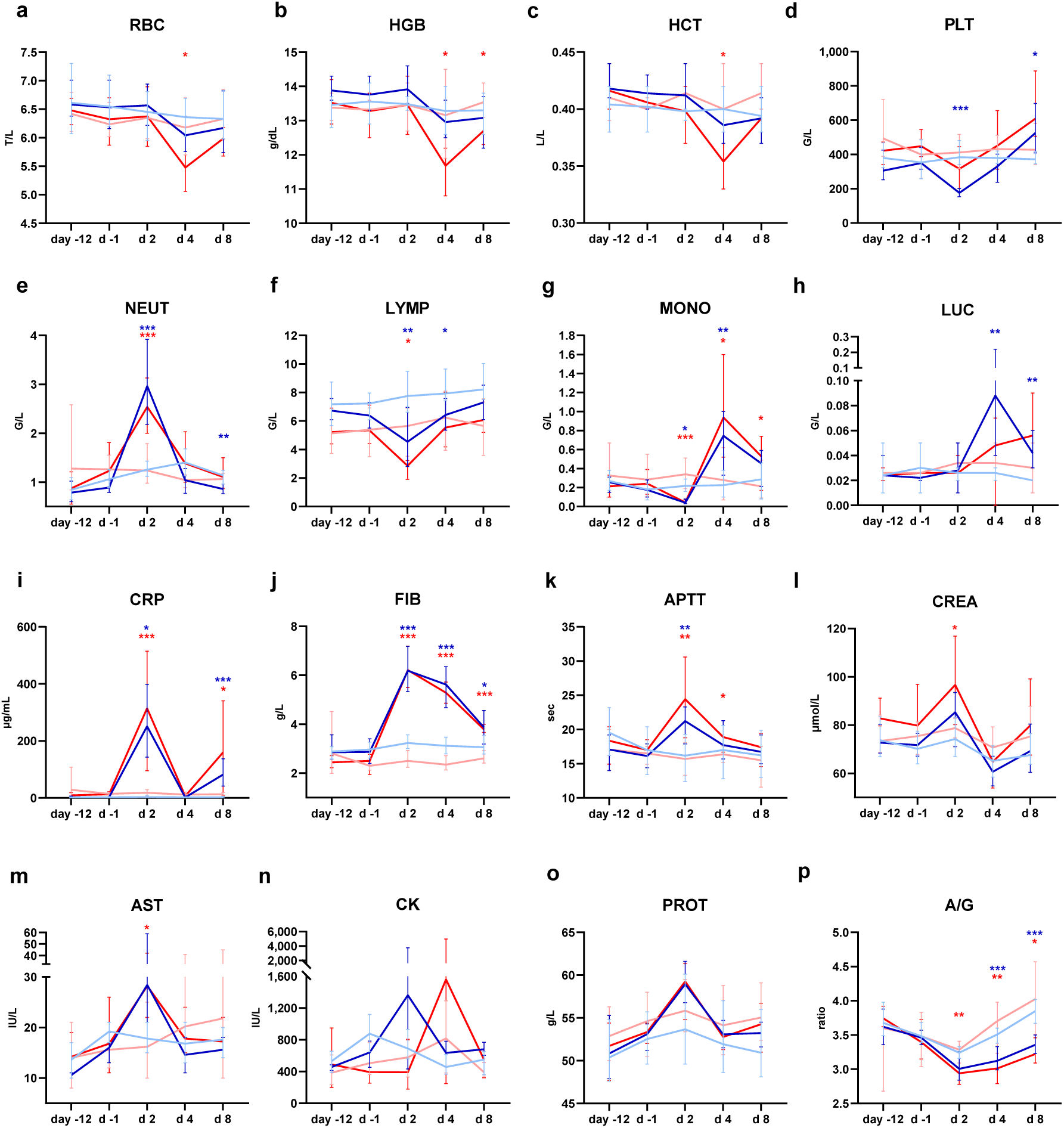
Hematology, coagulation and blood biochemistry in rabbits receiving a single saline or GRAd-COV2 administration. hematology parameters associated with main changes in GRAd-COV2 receiving rabbits compared to controls receiving saline are reported: (a) Red Blood Cell (RBC) counts, (b) Hemoglobin (HGB, grams per dL), (c) Hematocrit (HCT, liter of cells per liter of blood), (d) platelets (PLT), (e) Neutrophil (NEUT), (f) Lymphocyte (LYMP), (g) Monocyte and (h) Large Unstained Cell (LUC) counts. Counts are expressed as either Tera (T=1012) or Giga (G=109) per liter (L) of blood. Blood biochemistry and coagulation parameters associated to main changes: (i) C-Reactive Protein (CRP, μg/mL), (j) Fibrinogen (FIB, grams/liter), (k) Activated Partial Thromboplastin time (APTT, seconds), (l) Creatinine (CREA, μmol per liter), (m) Aspartate Aminotransferase (AST, International Units per liter), (n) Creatine Kinase (CK, International Units per liter), (o) total proteins (PROT, grams per liter) and (p) Albumin to Globulins ratio. Throughout, data represent the mean with range per study day of each treatment group, and the color code of the curves indicates treatment group assignment, either control male (light blue) and female (pink) animals, or GRAd-COV2 vaccinated male (blue) and female (red) animals. Statistically significant differences from controls of same sex, according to the statistical analysis decision tree, are indicated with blue (male) or red (female) asterisks: *(p=<0.05); **(p=<0.01); ***(p=<0.001)

Overall, both GRAd-COV2 and GRAdHIVNE1 vaccines were well tolerated; there were no mortalities, adverse clinical signs, local reaction at injection site and ophthalmological findings.

After the first vaccine administration, test item related in-life findings for GRAd-COV2 consisted of minor transient mean body weight (Fig. 2a) and food consumption Fig. 2b) reductions on the day following vaccination; this effect was more marked in the SD study, where a toxicology lot of the GRAd-COV2 vaccine was used, than in the RD study where animals were dosed with the clinical lot. GRAdHIVNE1 administration did not result in any significant body weight changes. However, GRAdHIVNE1 vaccination induced a transient decrease in food consumption in male rabbits after the first dose which returned to normal food consumption levels by day 6 and remained consistent throughout the study.

A minimal transient increase in mean body temperature (Fig.2c and Supplementary tables 4 and 14) 24h post first administration of both GRAd-based vaccines was recorded in all studies, with temperature equal or above 40°C in a subset of animals (3/5 female and 1/5 male in SD study, and 3/10 female in RD study for GRAd-COV2; 2/10 male and 8/10 female for GRAdHIVNE1), returning to normal by 48h. The second administration was better tolerated for both vaccines in RD studies, with no significant impact on body weight gain, food consumption or body temperature changes.

### GRAd vaccines induce transient non-adverse inflammatory changes in blood parameters

In general, clinical pathology parameters affected were suggestive of a systemic inflammatory response that was more pronounced following the first vaccination in comparison to the second one. An inflammatory leukogram (increased neutrophils, large unstained cells and monocyte counts, decreased lymphocytes, eosinophil and basophil counts), increase in acute phase protein concentrations, a slight decrease in platelets counts after three days from the first administration and a decrease in A/G ratio reflecting an increase in globulins concentration was observed, consistent with a transient systemic inflammatory reaction.

Hematology findings (Fig. 3 and Supplementary tables 5 and 15) consisted of transient decreases in red blood cell mass parameters (red blood cell counts, hemoglobin, hematocrit, Fig. 3 a to c) peaking at day 4 post administration. These changes in red blood cell mass parameters were noted in the GRAd-COV2 SD study and in the GRAdHIVNE1 RD study after the first administration, but not in the GRAd-COV2 RD study.

Decreased platelet counts (Fig. 3 d) were noted after the first GRAd-COV2 administration, but not after the second administration. Platelet counts were back to normal at day 4. Consistently, platelet counts were found in the normal range 4 days after GRAdHIVNE1 administration.

As for the leukocyte profile, 1 day post GRAd-COV2 administration, neutrophils (Fig. 3 e) counts increased, while lymphocytes (Fig. 3 f), eosinophils and basophils counts decreased, and all were back to baseline at day 8; after an initial drop at 1 day post administration, monocytes (Fig. 3 g) sharply increased at day 4, and were still elevated at day 8; LUC (Fig. 3 h) also increased at day 4 and were still elevated at day 8. All these changes resolved over time. Day 1 post first and second administrations were not sampled for GRAdHIVNE1, but the hematology observations at day 4 post administrations on red blood cell mass parameters and monocytes were consistent with those of GRAd-COV2.

A marked increase in C-Reactive Protein (CRP) and fibrinogen (Fig. 3 i-j, Supplementary tables 6-8 and 16-17) after each vaccination was noted for both vaccines, peaking 1 day post dose and gradually returning to baseline towards day 7. Other coagulation parameters were also transiently impacted: for GRAd-COV2, prolonged activated partial thromboplastin time (APTT, Fig. 3 k) was seen at day 2, returning to normal values thereafter. This prolongation was considered artifactual and secondary to the increase in CRP concentration, as it has been shown that increases in CRP concentration cause falsely prolonged APTT ^15^. For GRAdHIVNE1 study, where day 1 post dose was not sampled, APTT and PT were shortened 3 days after each administration compared to control groups, and were back to normal at end of study.

Other minor transient fluctuations (Fig.3 l to n, Supplementary tables 7 and 16) were noted in biochemistry parameters such as increased Creatinine, AST and Creatine Kinase; increased total plasma proteins concentration with a corresponding decrease of A/G ratio in absence of noticeable changes in albumin suggest an increase in the globulin component (Fig. 3 o-p), was detected in the GRAdHIVNE1 study. These changes were considered non-adverse and correlated to the inflammatory response associated with an immune response induction.

### Reversible macroscopic and microscopic changes following two repeated GRAd vaccine administrations

In rabbits necropsied after the acute phase, the mean weights of the spleen, the right iliac and popliteal lymph nodes, and the right inguinal lymph node (in males only) were moderately to markedly increased in animals of both sexes administered the vaccine (Supplementary table 9) when compared with control animals in the GRAd-COV2 RD study. The mean weights of the adrenal glands were slightly increased in males and the mean thymus weights were decreased in males and females administered with GRAd-COV2 (not statistically significant), suggestive of stress ^16^. In recovery phase animals, GRAd-COV2-related changes in organ weights were partially reversed in the draining lymph nodes (Supplementary table 10) and completely reversed for the spleen, adrenals and thymus.

GRAd-COV2-related macroscopic changes were limited to enlarged iliac lymph nodes in animals of both sexes necropsied after the acute phase, that completely resolved in the recovery phase animals (Supplementary tables 11-12). In general, microscopic changes consisted of partially reversible increased lymphoid cellularity in the spleen and lymph nodes draining the injection sites (Supplementary table 12), mostly characterized by an increased number of germinal centers (medulla to a lesser extent). In addition, reversible low-grade inflammation, mononuclear cell infiltration, necrosis, degeneration at the injection site and in tissues surrounding the injection sites were recorded in acute phase treated animals. Mononuclear infiltrates were composed of lymphocytes and plasma cells mainly. Mixed infiltrates were additionally composed of granulocytes. Other microscopic changes at the injection sites (myofiber degeneration/necrosis, myofiber regeneration, and hemorrhage) were also observed in both controls and vaccinated animals, after intramuscular injection that was attributed to the intramuscular injection procedure. Recovery animals showed complete reversion of microscopic changes at injection site (Supplementary table 13).

No GRAdHIVNE1 vaccine-related changes in terminal body weights, organ weights or organ weight ratios were identified, nor gross pathology lesions. GRAdHIVNE1 vaccine-related microscopic changes were confined to injection sites and adjacent skeletal muscle (Supplementary table 18). Microscopic changes in early euthanasia animals consisted of minimal to mild inflammation, necrosis, degeneration, and regeneration of skeletal muscle. Recovery animals at late euthanasia had fewer changes consisting of minimal to mild mononuclear cell inflammation in injection site and skeletal muscle suggesting partial resolution (Supplementary table 19).

These findings in the lymphoid organs and at the injection sites and surrounding tissues are expected changes following injection of a vaccine, and were not considered to be adverse.

### GRAd vaccines elicited potent immune response to the encoded transgene

For both GRAd-based vaccines, the induction of the expected specific adaptive immune response to the encoded transgene was assessed in blood samples collected 3 days post-second administration and at end of RD toxicity studies (recovery phase). For GRAd-COV2, high levels of antibodies to SARS-CoV-2 Spike were induced in treated animals only, and maintained until day 43 at endpoint titers above 1:100,000, as measured in serum of vaccinated animals by ELISA (Fig. 4 a-b). For GRAdHIVNE1, T cell responses to HIV networked epitopes (HIVNE) encoded in the transgene were clearly detected in peripheral blood mononuclear cells (PBMC) as measured by IFN-γ ELISpot. Such T cell response peaked 3 days after second vaccine administration, declining over time but still detectable at end of recovery phase in most animals (Fig. 4c).

**Figure 4.**
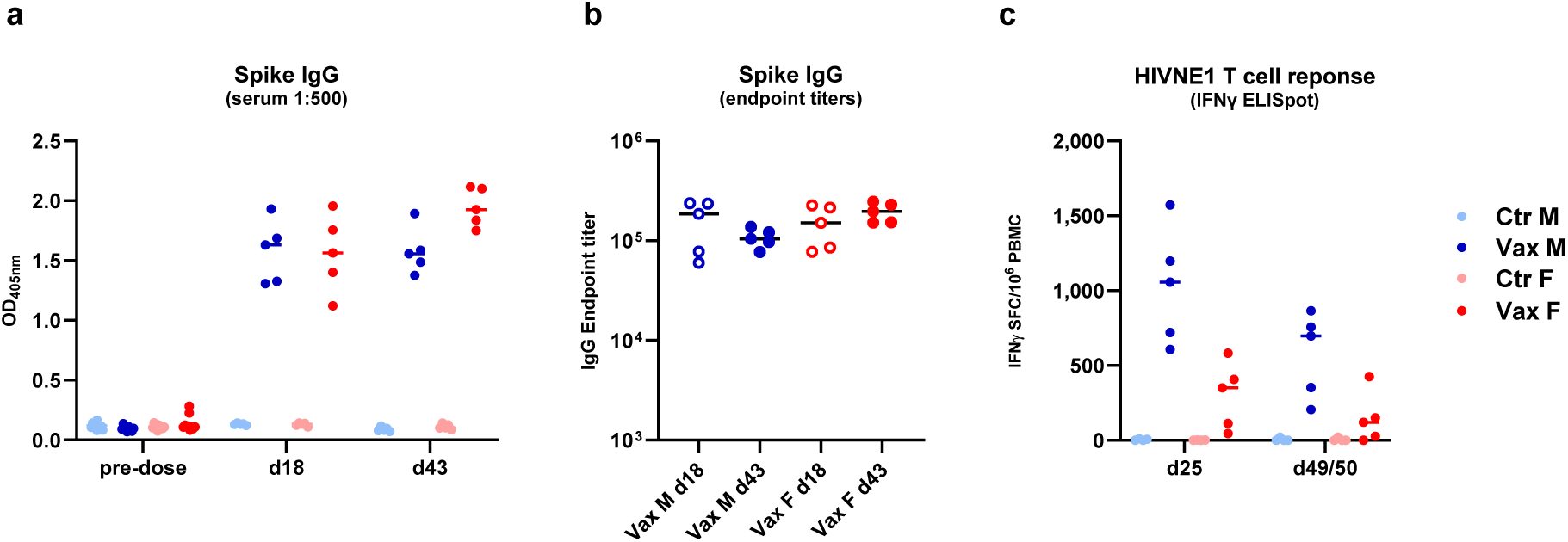
GRAd-COV2 and GRAdHIVNE1 immunogenicity. (a) Seroconversion to SARS-CoV-2 Spike in sera tested at 1:500 dilution in ELISA in rabbits from the GRAd-COV2 RD toxicity study. Each symbol represents antibody responses pre-dose or either at day 18 (3 days after 2nd GRAd-COV2 vaccination) or day 43 (4 weeks after 2nd vaccination). All data are expressed as optical densities (OD at 405 nm). (b) Endpoint IgG titers to Spike measured in sera from GRAd-COV2 vaccinated animals by ELISA with serum dilution curves. Each symbol represents the endpoint titers of sera from vaccinated male (blue symbols) and female (red symbols) animals at day 18 (empty symbols) and day 43 (filled symbols). (c) T cell response to the encoded HIVNE1 antigen measured by IFN-γ ELISpot in PBMC isolated from rabbits in the GRAdHIVNE1 RD toxicity study either at days 25 (3 days after 2nd GRAdHIVNE1 vaccination) or 49/50 (4 weeks after 2nd vaccination). All values are expressed as mean IFN-γ spot forming cells (SFC) per million PBMC. Throughout, the color code indicates treatment group assignment, either control male (light blue) and female (pink) animals, or vaccinated male (blue) and female (red) animals.

These data show that the repeated dose administration of GRAd-COV2 and GRAdHIVNE1 vaccines effectively triggers the immune system in New Zealand White rabbits and elicits the expected antibody or cell-mediated responses to the encoded antigens, validating the animal model used for safety assessment. The data also confirms that the rabbit is a less expensive, readily available, scientifically valid large animal model alternative to non-human primates for Ad-based vaccines like GRAdHIVNE1 that is specifically designed to induce a T cell response. The availability of well qualified methods for PBMC isolation and cryopreservation and high-quality rabbit-specific reagents to characterize cell-mediated immune response has made this upgrade practically feasible.

### GRAd vaccines share a common safety profile with non-replicating Ad vectors

A substantial amount of non-clinical safety data for intramuscular administration of vaccines based on replication defective Ad vectors has accrued over the years. A summary of most relevant published toxicity and biodistribution study designs can be found in Supplementary table 20. In general, a very predictable and favorable reactogenicity, safety and biodistribution/shedding profile emerges from these studies.

More specifically, a closer look to the most comparable toxicity studies in terms of animal model (rabbit), intensive sampling schedule and endpoints reveals that ChAd3-EBO-Z, a candidate vaccine for Ebolavirus based on the group C ChAd3 vector, was locally well tolerated and there was no evidence of systemic toxicity apart from minor transient increase in body temperature and decrease in food consumption and body weight ^17^. Reversible non-adverse changes in hematological and biochemical parameters, as well as mild inflammatory/degenerative changes at injection site and lymphoid compartment engagement were also observed in GRAd-based vaccines studies reported here The same profile emerged from the published toxicology package of ChAd155-RSV, a candidate vaccine for Respiratory Syncytial virus based on a different species C chimpanzee adenoviral vector ^18^. The findings were minimal in this study, possibly due to the lower dose injected. Very similar findings were described in a comprehensive paper describing a series of toxicology studies in rabbits of candidate vaccines based on adenoviral vectors of human origin (Ad5 and Ad35) encoding HIV-1, Ebola or Marburg antigens ^19^. Reported findings consisted mainly of transient blood cell population fluctuations (increased counts of neutrophils and LUC, decreased Lymphocytes, Eosinophils, Basophils, Platelets, initial drop and subsequent elevation of Monocytes; in some cases, reductions in Hemoglobin/Hematocrit), marked transient increases of acute phase proteins (mainly Fibrinogen, CRP and globulins), prolonged APTT, shortened PT and other more sporadic observations. This pattern was considered non-adverse and consistent with inflammatory leukogram, transient inflammation and induction of an immune response to vaccination. Even if less information is available for non-clinical studies of two of the Ad vectored vaccines approved for COVID-19 ^20,21^ the overall emerging local and systemic toxicity profile and microscopic findings were quite similar.

The GRAd-COV2 and GRAdHIVNE1 biodistribution profiles are aligned with published experience for human and simian adenoviral vectors injected intramuscularly in Rabbits, Rats or Mice, as reviewed and summarized in Table 2. Such studies all showed that adenoviral genomes were found mainly at the injection site and draining lymph nodes, persisting for weeks with steadily decreasing levels over time, indicating clearance. Genomes were not quantified in brain, heart, kidney, lung, and most relevantly in ovary and testis samples. Sporadic and transient (mostly 24 hours only post-injection) exposure of blood, spleen, and liver were observed in all studies. Importantly, the Ad genome quantities noted at study termination in both muscle and lymph node stations were in all cases <30,000 copies/μg of host cell DNA, the threshold above which further DNA integration studies would be warranted ^22^. A shedding study of ChAd155-RSV administered intramuscularly in rats ^18^ showed that genomes were generally not detectable in any secretions or excreta samples. The described biodistribution/shedding profile was mostly confined locally and is consistent with the replication defective nature of all tested vectors, enabling only one round of cell infection, with no further amplification and dissemination of viral progeny.

**Table 2:**
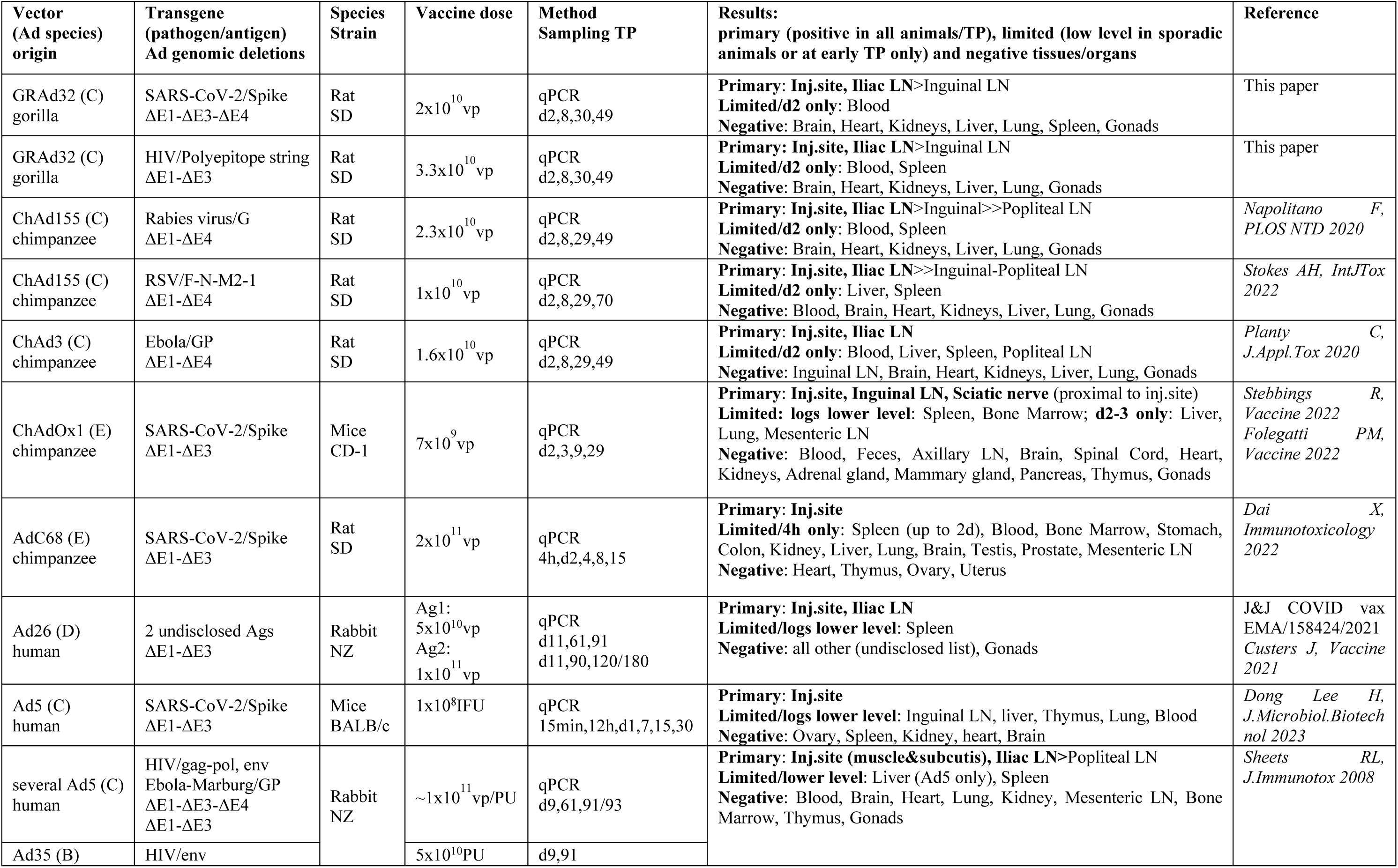
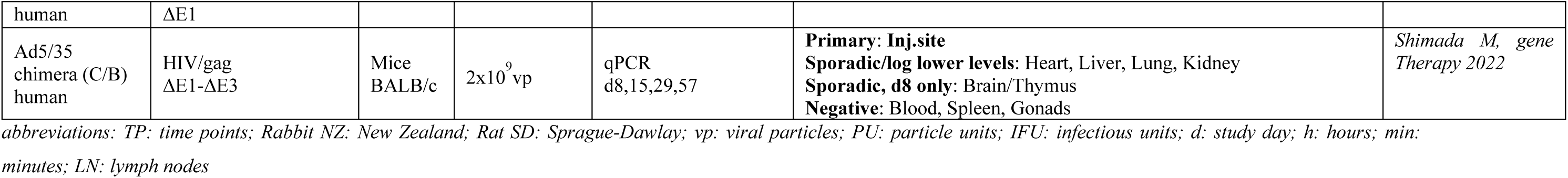
Biodistribution studies of replication deficient adenoviral vectors by intramuscular, single administration.

In summary, all non-clinical safety data suggest that the biodistribution and toxicity profile of various replication-defective Ad vectors administered intramuscularly is mostly dictated by the technology platform and to a very limited extent by the encoded transgene.

## Discussion

Multiple toxicity and biodistribution studies were completed as part of the pre-clinical evaluation for two vaccines based on the GRAd vector platform. The consistently acceptable safety profile demonstrated by both these vaccine candidates ensured regulatory approval for the initiation of their respective clinical trials. We confirm here that GRAd-COV2 and GRAdHIVNE1 non-clinical safety profiles are very well aligned with our previous experience with simian non-replicative adenoviral vectors ^17,18,23^, and with similar published studies of other simian ^24,25^ and human-derived adenoviral vectors (^19,26 20^), including the widely used COVID-19 vaccines from AstraZeneca and Janssen ^20,21^. The safety, immunogenicity, and efficacy of the Ad vaccine platform in humans has been validated in multiple clinical studies and in clinical practice for Ad-based COVID-19 vaccines during the last decades. The GRAd-COV2 phase 1 and phase 2 clinical trials were conducted cumulatively in ∼700 participants ^12,13^, consisting of healthy young and elderly cohorts and subjects with co-morbidities, inclusive of a small subset of people living with HIV (PLWH). The safety and reactogenicity profiles were the same as those observed in clinical trials of other Ad based vaccines in terms of type, severity, and duration of local and systemic adverse events.

The findings associated with intramuscularly administered adenovirus-based vaccines, including systemic adverse events (AEs) and changes in clinical pathology parameters, typically occur within the first 24-72 hours after the injection. This timing is noted in both non-clinical toxicology studies and in clinical trials, as well as in real-world experience with approved Ad-based COVID-19 vaccines. It aligns with the body’s innate immune response recognizing the “viral nature” of the vaccine’s vector component and of the expressed transgene, which in mouse studies is detectable at the mRNA level at the administration site for up to 2 weeks ^27,28^. Blood transcriptomics studies in humans concur with this figure, showing a sharp increase in inflammatory and antiviral gene expression pathways, particularly those dependent on Interferon, within a few hours to a day after Ad vaccination ^29,30^. These levels usually return to normal within a week. Indeed, vaccines based on Ad vectors are considered “self-adjuvanting” and do not require or benefit from the co-administration of classical vaccine adjuvants. Instead, the capacity of the Ad vectors to activate innate immunity through a variety of Pathogen-Associated Molecular Patterns (PAMP, i.e., capsid proteins and nucleic acids) that upon administration induce pattern-recognition receptor activation, immune-cell recruitment to the injection site and draining lymph nodes, and cytokine production ^31^ is what shapes the adaptive immune response to the transgene. The adaptive immune response to the vaccine encoded antigen, whether it stimulates antibodies or T-cells, or both, typically becomes detectable between 1-2 weeks after vaccination and peaks around a month post administration, at which point the Ad vaccine reactogenicity is no longer observed.

The type of antigen encoded in Ad-based vaccines does not seem to play a major role in their safety or biodistribution profile, even though some differential transcriptomic response in infected cells depending on differential TLR triggered pathways upon transgene expression may be expected and have been described in mice ^32^. Similar reactogenicity and safety profiles have been observed in the reported toxicology studies, whether the Ad vectors encode traditional antibody-inducing antigens (like full-length viral glycoproteins from Ebola, Rabies, or RSV) or transgenes designed to trigger T-cell responses only. Relevantly for GRAdHIVNE1, there are no specific safety concerns associated with the novel HIV transgene design aimed at inducing CD8^+^/Th1 T cell response, since polyepitopic strings encoded by viral vectors are in clinical trials for infectious indications like HIV ^33^ and cancer ^34,35^, with no related safety signals reported.

Of note, the Ad-based vaccines showing similar safety profiles in the non-clinical studies considered here originated from different manufacturers, were produced in different cell substrates, and carried different genomic deletions in the adenovirus early genes (ΔE1, ΔE1-ΔE3, ΔE1-ΔE3-ΔE4, ΔE1-ΔE4, supplementary table 20), again pointing toward the applicability of a platform approach.

Replication-defective Ad vectors can only undergo one single round of cell infection upon administration and there is no secondary dissemination of viral progeny. Their biodistribution is expected to depend on the administration route and on the Ad capsid proteins (i.e., the specific Ad vector serotype/isolate and their interaction with cellular receptors) without any influence of the antigenic cargo. In contrast to other vaccine platforms like VSV, where the capsid is pseudotyped with the target vaccine glycoprotein, the encoded antigens are not incorporated in the Ad viral particles and do not therefore alter their tropism. When transgene expression-repression systems are adopted in the Ad vector platform for improving vaccine productivity, as is the case for all GRAd-based vaccines and the production cell substrate (ReiCells35S), the encoded antigen is virtually not expressed and thus not co-purified with the vaccine particles during the manufacturing process. However, in published studies the reported biodistribution is similar not only within species C and E (human or simian) adenoviral vectors like GRAd, sharing CAR as their primary cellular receptor, but also for Ad35, a species B vector which utilizes CD46 as cellular receptor, and for Ad26, a species D vector whose primary receptor usage is debated among CD46, CAR, and sialic acid-bearing glycans. Thus, upon intramuscular administration the distribution of the vaccine inoculum is mostly confined to the muscle and surrounding tissues and to the draining lymphatic compartment, and the potential Ad serotype tropism is less relevant than when administration occurs through intravenous route. As for all other Ad vectors considered here and according to the well-established non-integrating nature of adenoviruses, GRAd-COV2 and GRAdHIVNE1 genome quantities noted at biodistribution study termination were <30,000 copies/μg of host cell DNA in both injection site (muscle) and draining lymph nodes, the threshold above which further DNA integration studies would be warranted ^22^.

The relevant toxicity studies have been accurate in predicting the clinical safety profile of Ad-based vaccines, consisting in mild to moderate inflammatory local and systemic effects like mild pyrexia and minor transient changes in blood parameters. However, extremely rare AEs like the Vaccination-induced thrombosis with thrombocytopenia (VITT) that have been described for AstraZeneca and Janssen COVID-19 vaccines during their use in mass vaccination campaigns cannot be predicted by non-clinical studies. VITT was not identified in any of the studies conducted with the two vaccines, which demonstrates a limitation of the animal model in predicting this clinical liability. There is still lack of clarity on the condition’s underlying mechanism(s) and therefore an appropriate animal model is yet to be identified, and it is yet unknown if the syndrome may be related or not to SARS-CoV-2 Spike antigen only, or to aberrant Spike/Ad proteins’ splice products. Although the molecular basis of VITT remains unclear, vector genomic configuration may play a role in modulating splicing artifacts. Relevant to this point, the transgene expression cassette in the GRAd vector is inserted in the deleted E1 region, close to the left ITR of the viral genome, as per standard practice. However, differently from ChAdOx1 nCoV-19 and Ad26.COV2.S, the direction of transcription of the expression cassette in GRAd is pointed towards the left ITR. This orientation virtually abolishes the possibility of generating aberrant/chimeric transgene/Ad protein products *in vivo* due to alternative splicing events ^36^, a feature that may be seen as an additional safety attribute for the GRAd vaccine platform.

Taken together these observations suggest that the biodistribution and the toxicity profiles are common to the vaccine platform (i.e. replication-defective Ad vector of any derivation) with no significant influence of the encoded transgene. Our non-clinical package therefore functions as “platform evidence” for GRAd technology when the vector, route and dose range are conserved, and the transgene product lacks known or suspected toxicity. These convergent profiles are the type of knowledge envisaged by the FDA’s Platform Technology Designation (PTD) framework for drug development ^37^. This framework allows evidence generated (“prior knowledge”) on a characterized platform to inform the development of subsequent products that share core technological elements. In parallel, Europe is exploring the concept of a “master file” for vaccine platform technology. The EMA has recently implemented this approach for the veterinary vaccine sector ^38^, signaling momentum toward the structured reuse of platform data.

According to main WHO, EMA and FDA regulatory guidelines, a repeated-dose toxicity study with local tolerance and a bio-distribution study are recommended for progressing any novel vaccine candidates into Phase I trials. Nevertheless, recently during the COVID-19 pandemic and Ebola outbreaks there has been increased consideration and guidance with WHO and Regulatory Authorities on a vaccine platform approach for non-clinical safety and quality aspects, with the intent to accelerate the process of research and development as well as the global mobilization of vaccine interventions for addressing infectious diseases that threaten Global Health ^39–41^. A vaccine platform approach that can streamline or modify the standard requirements for advancing novel candidates into clinical studies is particularly valuable during epidemics and pandemics, when speed is critical, as exemplified by initiatives such as CEPI’s “100 Days Mission.” ^42,43^. Such an approach is equally important in complex vaccine development contexts, such as HIV, where numerous innovative platform-based candidates are being explored to address the intrinsic biological complexity of the virus, which has long hindered the development of an effective vaccine. An experimental medicine framework is envisioned, where multiple iterations/combinations of vaccine candidates need to be rapidly tested and down-selected in early-phase clinical trials, to accelerate the optimization of much needed efficacious vaccines for Global Health ^8^. Viruses that cause some diseases such as influenza and COVID-19 are subject to rapid genetic evolution resulting in vaccines being less effective. In these cases, rapid development of vaccines with modified antigenic content is necessary to prevent and control outbreaks. Inactivated influenza vaccines have for many decades used a platform technology of vaccine subunits to yearly deliver the updated virus antigens, leveraging on standard manufacturing process and formulation. Similarly, for COVID-19 mRNA vaccines, a platform approach is currently employed, to rapidly incorporate the Spike mRNA from the circulating SARS-CoV-2 strain into an updated, strain specific vaccine. In both cases, development timelines are substantially shortened by eliminating the need for pre-clinical safety studies in laboratory animals and extensive clinical studies prior to general use in the community.

Based on the non-clinical safety database generated for two GRAd-based vaccines, together with published pre-clinical and clinical data from other adenoviral-vectored vaccines and evidence from clinical trials demonstrating the robust safety of the GRAd-COV2 vaccine in humans, we propose that sufficient scientific evidence exists to justify deferring additional non-clinical evaluations for new GRAd-based vaccine candidates intended for intramuscular administration, in alignment with a vaccine platform approach. This approach is reasonable for any Ad vector for which non-clinical and clinical evidence has already been already generated with at least one transgene, provided that the new antigen is free of known or suspected toxicity; however, a change in the intended administration route or modifications to vector capsid proteins that may impact tropism would require a full non-clinical evaluation as per standard practice. A comparable regulatory approach, grounded in similar observations and considerations, has recently been proposed for non-clinical safety of vaccines formulated with the AS01 adjuvant ^44^.

This strategic approach can accelerate vaccine development timelines for critical diseases affecting humanity. Furthermore, reducing the use of laboratory animals, especially when formal non-clinical studies may ultimately prove redundant and the added value of data is limited ^45^, aligns perfectly with the 3R principles (Replacement, Reduction, Refinement). These findings, therefore, reinforce the suitability of the GRAd platform for streamlined non-clinical assessment and exemplify a data-driven application of the 3R’s in vaccine development.

## Methods

### Test and Control Items

The vaccines’ stock concentration and dosage are expressed in viral particles (vp) as measured by qPCR, representing encapsidated GRAd genomes. The GRAd-COV2 toxicology lot used for the single-dose toxicity study, was provided frozen in 2 mL polypropylene cryovials containing 1mL solution at 2 × 10^11^ vp/mL, while the GRAd-COV2 clinical lot, used for the repeated-dose toxicity and the biodistribution studies, was provided frozen in operculated transparent glass vials filled with 1.2 mL solution at 2 × 10^11^ vp/ml. The GRAdHIVNE1 toxicology lot, used for repeated dose and biodistribution studies, was provided frozen in 4.5 mL polypropylene cryovials containing 3 mL solution at 3.3 × 10 ^11^ vp/mL. GRAd vaccines were formulated in A195 buffer: 10 mM TRIS, 10 mM histidine, 75 mM NaCl, 5% sucrose, 1 mM MgCl2, 0.02% (w/v) polysorbate-80, 0.1 mM EDTA, 0.5% ethanol (w/v), pH 7.4. All vaccines were stored at −80°C until use. The control item was a ready-to-use nonpyrogenic, sterile 0.9% saline formulation.

### Animal Welfare

The two biodistribution studies and the GRAd-COV2 toxicity studies were carried out at Charles River Laboratories (CRL) Evreux (France), a French National Authority certified Test Facility. The studies were conducted in compliance with the General Principles Governing the Use of Animals in Experiments (Council Directive 2010/63/EU of September 2010 and French decree No. 2013-118 of 01 of February 2013) and following the general procedures for animal care and housing AAALAC International recommendations. The CRL Evreux Ethics Committee (CEC) reviewed the study plans in order to assess compliance with the corresponding authorized “project” as defined in Directive 2010/63/EU and in French decree No. 2013-118. The GRAdHIVNE1 toxicity study was carried out at AmplifyBio Ohio (USA). The study was conducted in compliance with the general procedures for animal care and housing meeting AAALAC International recommendations and the current requirements stated in the “Guide for Care and Use of Laboratory Animals” (National Research Council, Current Edition), and conformed with the testing facility SOP. The study protocol was reviewed and approved by the Institutional Animal Care and Use Committee (IACUC). Veterinary care was available throughout the course of all studies.

### Biodistribution studies design, clinical and pathology examination

Two biodistribution studies were conducted in Sprague-Dawley (Crl:CD(SD)) rats with the GRAd-COV2 and GRAdHIVNE1 vaccines. Both studies followed established practices and standard operating procedures that are compliant with the OECD Principles of Good Laboratory Practice, but GLP status was not claimed.

Male and female Sprague-Dawley (Crl:CD(SD)) rats were obtained from Charles River Laboratories and acclimated to the study conditions for at least 7 to 11 days prior to the start of the study. On the first day of treatment, the animals were 8 to 9 weeks old. The males weighed 280-467 g and the females weighed 189-273 g. They were group housed in polycarbonate cages (n = up to 5 animals/sex/group) containing autoclaved saw-dust. Both the biodistribution studies envisaged a control group, and a vaccine administered group, housed in separate racks in rooms each with filtered air (8-15 cycles/hour of filtered, non-recycled air) at a temperature within the range 20-24°C and relative humidity within the range 30%-70%. The lighting followed a 12-hour light/12-hour dark cycle. Rats had free access to a standard laboratory rat diet (SSNIFF A04 Mod. Maintenance pelleted diet, sterilized by irradiation) and to 0.22-μm filtered drinking water. For psychological/environmental enrichment, animals were provided with at least two items (e.g. hut, nylabone, tunnel), except when interrupted by study procedures/activities.

Rats in the vaccinated group were randomly allocated to four sub-groups (one for each time point) of 10 animals, each comprising five males and five females. These animals received a single intramuscular injection (100 μL or 200 μL) of the investigation vaccine at a dose of 2 × 10^10^ and 3.3 × 10^10^ vp/injection of the GRAd-COV2 and the GRAdHIVNE1 vaccines respectively in the right quadriceps. The dose of vaccines administered in the two biodistribution studies approximates 1/10th of the proposed human clinical dose, assuring a 20 to 30-fold higher margin of exposure in rats compared to the proposed human clinical dose on a per kilogram body weight basis. A sub-cohort of vaccinated rats were necropsied 24 hours 7, 28-29 or 48 days after vaccination (i.e., days 2, 8, 29-30 and 49, respectively). Sham vaccinated control animals (four males and four females) were administered sterile saline (0.9% NaCl) for injection. One rat/sex was necropsied at the same time point as in the treated group.

Study animals were checked for mortality at least once a day during the acclimation period and twice a day during the treatment and observation periods. Each animal was observed once a day before the beginning of the treatment period, on the treatment day and until the end of the study and daily cage-side observation was performed at approximately 1 and 6 hours postdosing. In addition, detailed clinical examinations were performed on all animals once before the beginning of the treatment period, then once a week until the end of the study. The body weight of each animal was recorded once during the pre-treatment period, on the day of vaccination (before injection), once a week during the observation period and on the day of euthanasia. The quantity of food consumed by the animals in each cage was recorded once a week from the first day of treatment until the end of the study. On completion of the observation period, the animals were euthanized by an intraperitoneal injection of sodium pentobarbital followed by exsanguination, and a full macroscopic post-mortem examination was performed. Designated organs were weighed and selected tissue specimens collected to quantitate the genome copy biodistribution by quantitative polymerase chain reaction (qPCR), as described below.

### Organs and tissue sample collection, processing and DNA extraction

Blood was collected before necropsy from the abdominal aorta or vena cava of animals (under deep anesthesia) into potassium EDTA tubes. Blood was aliquoted, snap frozen in liquid nitrogen and then stored at −80°C until DNA extraction. All tissues from control group animals were collected and extracted in a dedicated room to avoid cross-contamination. For vaccinated animals, a specific sequence of sample collection, processing and DNA extraction from tissues was followed. Tissues with potential for lower genome copy (gc) titers were collected first followed by the ones with potential for higher gc titers.

DNA was extracted from tissue samples using the NucleoSpin Tissue kit (Macherey-Nagel) and from blood samples using the NucleoSpin Blood QuickPure kit (Macherey-Nagel). At the end of the extraction process, DNA was eluted in elution buffer BE (i.e. 70 µL for blood samples and 100 µL for tissue samples).Sentinel control samples were included within each DNA extraction batch to monitor the level of potential cross-contamination between samples during the extraction and PCR processes.

### Biodistribution evaluation by quantitative polymerase chain reaction

The qPCR method was originally validated for GRAd-COV2, and subsequently re-qualified for GRAdHIVNE1 by CRL Evreux. The GRAd-COV2 genomic DNA was used as the analytical standard for the PCR assay used for sample analysis from both biodistribution studies. The GRAd-COV2 and GRAdHIVNE1 genomic DNA share a common CMV promoter region upstream of the transgene, and the qPCR assay primers and probe are located in this common region of the vector. Primers and probe were procured from Integrated DNA Technologies (IDT). The details are as follows: forward primer CMV: 5’-CATCTACGTATTAGTCATCGCTATTACCA -3’; reverse primer CMV: 5’-GACTTGGAAATCCCCGTGAGT -3’; Taqman Probe CMV: 5’-56FAM ACATCAATG/ZEN/GGCGTGGATAGCGGTT/3IABkFQ -3’. Based on method qualification studies, the primers and probe final concentrations were set at 200 nM and 400 nM, respectively. The qPCR reaction mixes (50 µL) contained 10 µL of DNA sample (reference analyte, control or in-study sample DNA, 1 μg whenever possible) and 40 µL of a Taqman Gene Expression Master Mix (Applied Biosystems). qPCR plates were run using a QuantStudio 7 instrument (Applied Biosystems). Amplification was detected in real-time over 40 cycles (including an elongation step at 60°C) by following the evolution of the fluorescent signal generated by the degradation of the probe. All samples were analyzed in duplicate, in parallel with a calibration curve containing eight calibration standards and two sets of QC samples at three levels prepared by spiking GRAd-COV2 genomic DNA in 1 µg of Herring Sperm DNA. Corresponding Ct values were plotted as a function of the base ten logarithm of the viral genomic DNA quantity of calibration standards, and the resulting linear curve was fitted using a validated Microsoft Excel sheet. GRAd-COV2 and GRAdHIVNE1 DNA quantity in samples was then interpolated from the calibration curve. For all samples, DNA quantity was calculated by interpolation from the calibration curve and was expressed as genome copy numbers per µg of DNA.

### GRAd-COV2 and GRAdHIVNE1 toxicity evaluation in rabbits

The New Zealand White rabbit model was chosen for the toxicity evaluation based on recommendations by EMA and WHO for this type of investigation (EMA-CPMP, 1997; WHO, 2003), and on previous experience. The relevance of the animal model used for the repeat dose toxicity study was validated by the demonstration of specific antibody or cell mediated response induced by the transgene expressed in the GRAd-COV2 and GRAdHIVNE1 vaccines respectively. Supplementary Table 1 highlights differences and commonalities in the three study designs.

### GRAd-COV2 single dose toxicity study

The objective of the single dose (SD) toxicity study was to evaluate the local tolerance and systemic toxicity of the GRAd-COV2 vaccine, after a single intramuscular administration to rabbits followed by a 7-day observation period. The GLP study was conducted at CRL Evreux, France.

Two groups of New Zealand White rabbits, each comprising of five animals/sex received either sterile 0.9% saline or GRAd-COV2 at 1 × 10^11^ vp/animal once by intramuscular route under a constant dosage volume of 500 µL/animal followed by a 7-day observation period. Injections were done in the anterior thigh muscle. Multiple parameters were evaluated to assess the toxicity of the investigational vaccine. Mortality and morbidity check were done at least once a day during the acclimation period and at least twice a day during the treatment and observation periods. Cage side observation was done at least once a day for identify clinical signs. Detailed clinical evaluation was done pre-vaccination, on day of treatment and on days 4 and 7. Local injection site reactivity was evaluated by Draize scoring pre-vaccination, and after 4-, 24-, 48- and 72-hours post vaccination. Body weight and food consumption was recorded pre-dose, on day of treatment and once a day till end of the study. Body temperature was monitored pre-vaccination, and at 4- and 24-hours, post vaccination. Blood was sampled from the rabbits for clinical pathology evaluation (hematology, clinical chemistry, coagulation and c-reactive protein) pre-dose, and on days 2, 4 after vaccination and at terminal necropsy on day 8.

For hematology parameters, blood was collected into potassium EDTA tubes, and the blood composition was analyzed using the ADVIA 120 Hematology Analyzer. The parameters measured were erythrocytes, mean cell volume, packed cell volume, hemoglobin, mean cell hemoglobin concentration, mean cell hemoglobin, thrombocytes, leucocytes and the differential white cell count with cell morphology and reticulocytes. Blood smears were microscopically examined if the blood sample was not accepted by the analyzer. Microscopic evaluation was also systematically performed for the identification and gradation of platelet aggregates when platelet count was <100 G/L. Blood was also collected into sodium citrate tubes and analyzed using ACL TOP 550 CTS blood coagulation analyzer for the determination of prothrombin time (PT), fibrinogen level and activated partial thromboplastin time (APTT). For clinical chemistry analyses, blood was taken into heparinized tubes and analyzed using the ADVIA 1800 blood biochemistry analyzer. The parameters measured were alkaline phosphatase, alanine aminotransferase, aspartate aminotransferase, creatine kinase, glucose, total protein, albumin (and the albumin/globulin ratio), urea, creatinine, bilirubin, cholesterol, triglycerides, calcium, sodium, potassium, chloride and inorganic phosphate. C-Reactive Protein (CRP) levels were determined in each animal, in pre-dose, and on days 2 and 4 after vaccination and at terminal necropsy on day 8 using a validated ELISA method. At the end of the study both control and vaccinated group of animals were euthanized by an intravenous injection of sodium pentobarbital followed by exsanguination, and tissues/organs were collected. The brain, draining lymph nodes (popliteal, inguinal and iliac), heart, injection site, kidney, liver and lungs with bronchi were collected and organ weights recorded. These tissues/organs were preserved in formalin. Only the draining lymph node and injection site were evaluated microscopically after H&E staining.

### GRAd-COV2 and GRAdHIVNE1 repeat dose toxicity study in rabbits

The dosing schedule was either N or N+1 where “N” refers to proposed number of clinical doses plus one additional dose. New Zealand White Rabbits were randomly allocated into two groups, a control and a treatment group in each of the repeat dose (RD) toxicity study. In the GRAd-COV2 RD toxicity study, conducted at CRL, Evreux, France, two groups of rabbits, each comprising of ten animals/sex received either sterile 0.9% saline or GRAd-COV2 at 1 × 10^11^ vp/animal twice by intramuscular injection at 2-weeks interval (days 1 and 15). Each of the rabbits on study were administered a constant dosage volume of 500 µL/animal intramuscularly to achieve the nominal dose. Injections were done in the right anterior thigh muscle. A subset of five rabbits/sex/group (main study animals) were euthanized 3 days after the second vaccination to evaluate the acute response post vaccination. The remaining animals (recovery phase animals) were kept for a 4-week treatment-free period to evaluate any delayed onset toxicity, identify resolution or aggravation of any acute toxicity findings.

The repeat dose toxicity study done with GRAdHIVNE1 vaccine in rabbits at AmplifyBio, OH, USA was almost identical to the GRAd-COV2 vaccine repeat dose tox study, except that the rabbits received the two vaccinations at three-week (day 1 and 22) dosing interval, the dose volume administered was reduced to 300 µL/animal due to the higher nominal titer of the vaccine and the number of rabbits allocated to the control group was limited to 8 animals/sex/group

### In-life observation, clinical and hematological parameters

Morbidity and mortality check by cage-side observation was done twice a day throughout the study. Detailed clinical evaluation was conducted pre-dose, on day of dosing (either 4 hours or at 1,6 and 24 hours) and once weekly thereafter for the GRAd-COV2 and GRAdHIVNE1 vaccines respectively. Draize scoring for local reactions at the injection sites were recorded pre-vaccination at 4, 24, 48 and 72 hours after each injection for the GRAd-COV2 vaccine. Draize scoring for local reactivity at injection site was evaluated pre-dose and post vaccination at 6, 24, 48 and 72 hours after each vaccination and at terminal necropsy in the GRAdHIVNE1 vaccine toxicity study. The body weight of each animal was recorded once prior to vaccination, then daily for 3 consecutive days post vaccination and subsequently once a week thereafter for the GRAd-COV2 RD tox study. In the GRAdHIVNE1 RD toxicity study the body weight was evaluated once prior to vaccination and post vaccination on days 2, 3, 8, 11, 15, 18, 21, 23, and 24 and twice weekly thereafter. Terminal body weight at necropsy was also recorded. The quantity of food consumed by the animals was recorded pre-dose, on day of dosing and then twice a week in the GRAd-COV2 toxicity study, while it was recorded daily in the GRAdHIVNE1 RD tox study. Food consumption was calculated per animal and per day. In the RD toxicity study with GRAd-COV2 rectal temperature was measured, once pre-vaccination, 4 and 24 hours after each administration. In the GRAdHIVNE1 toxicity study interscapular temperature of animals recorded using implantable programmable temperature transponder on both vaccination days at 6, 24, 48 and 72 hours post dose. Ophthalmological examinations were performed Pre-vaccination, and once before the main and recovery animals were euthanized.

Clinical pathology evaluation included hematology, clinical chemistry, coagulation and CRP evaluation. In the GRAd-COV2 repeat dose toxicity study blood sampling for clinical pathology evaluation were performed from recovery animals, once before the vaccination and a select set of clinical pathology parameters on multiple days post vaccination (days 2, 4, 8, 14, 16, 18). In addition, clinical pathology samples were collected from recovery animals on days 22 and 43. In the GRAdHIVNE1 vaccine RD toxicity study blood was collected on day 4 and at terminal necropsy from both main (day 25) and recovery group (day 50) of animals for evaluation of hematology, clinical chemistry and coagulation. For CRP levels only, blood was collected also pre-vaccination and on day 2 post vaccination.

For GRAd-COV2 RD study, the hematology, coagulation and biochemistry parameters as well as equipment and methods are the same described in detail for GRAd-COV2 SD study. For GRAdHIVNE1 study, Advia 2120i Hematology Analyzer, Stago Compact Max Coagulation Analyzer and Roche Cobas c501 Chemistry Analyzer were respectively used. The biochemistry parameters evaluated, in addition to those described for GRAd-COV2, also included gamma-glutamyltransferase, globulins, lactate dehydrogenase, hemolysis/Icterus/Lipemic indexes. A validated ELISA method was used for evaluation of the CRP levels for GRAd-COV2 study, while for GRAdHIVNE1 an immunoturbidimetric method on the Roche cobas platform was used.

### Necropsy, tissue processing and histopathological examination

In each of the repeat dose toxicity study, rabbits were deeply anesthetized by an intravenous injection of sodium pentobarbital and then euthanized by exsanguination. They were examined for external changes and gross pathological changes. Special attention was paid to the macroscopic examination of the injection sites.

Adrenals, brain, lymph nodes, epididymides, heart, kidneys, liver, ovaries (females), pituitary gland, prostate (males), spleen, testes (males), thymus, thyroid glands with parathyroid glands and uterus (females) were weighed and subsequently preserved in phosphate-buffered neutral 10% formaldehyde for microscopic evaluation. Other organs were not weighed but were preserved for examination, including aorta, colon, eyes and optic nerves (fixed in modified Davidson’s fixative), femoral bone, gall bladder, gut-associated lymphoid tissue, larynx, lungs with bronchi, pancreas, salivary glands, sciatic nerve, tri-ceps muscle, spinal cord, sternum with bone marrow, stomach, urinary bladder and vagina (females). All tissues submitted for histopathological evaluation were embedded in paraffin wax, sectioned at 4 μm and stained with hematoxylin and eosin.

To analyze the injection sites, the complete muscles were collected and preserved in phosphate-buffered neutral 10% formaldehyde. Three approximately 5-mm pieces from around the injection site were prepared, examined macroscopically for gross findings and processed for microscopic evaluation in three semi-serial sections, 150 μm apart, for each piece.

### Immunogenicity

In the GRAd-COV2 RD toxicity study, antibody response to the spike protein expressed was evaluated on blood samples collected prior to vaccination, on day 18 from all animals after the second vaccination and on day 43 at terminal necropsy of recovery animals, by means of a noncommercial SARS-COV2 ELISA assay. Briefly, HIS-tagged SARS-CoV-2 full length spike protein (Sino Biological) diluted to 1 μg/mL in assay buffer (PBS 0.2%BSA) was immobilized on Ni-NTA HisSorb plates (Quiagen) for 1h at 25°C. Serially diluted test sera were added to the coated plates and incubated for 2 hours. Binding of spike specific antibodies was detected using alkaline phosphatase conjugated anti-rabbit total IgG (Sigma) and p-Nitrophenyl Phosphate substrate (SigmaFast, Sigma). Absorbance was read at 405 nm using EnSight™ multimode microplate reader (Perkin-Elmer), and at 620 nm to remove the background. To assess seroconversion, sera samples were tested at a single dilution (1:500) in duplicate. A 6 point, 3-fold serial dilution curve was generated to determine the aniti-SARS-CoV-2 Spike antibody titers. Endpoint titers were calculated as the reciprocal of the highest serum dilution resulting in an optical density greater that 4-fold the control wells with secondary antibody alone.

In GRAdHIVNE1 RD toxicity study, peripheral blood mononuclear cells (PBMC) samples were isolated from blood samples collected from vaccinated and control animals on days 25 and 49/50. The T cell response was determined by the rabbit IFN-γ ELISpot plus kit from Mabtech and using the Millipore 96-filtration plates (MSIPS4W10). PBMCs (2 × 10^5^ per well) were plated and incubated for 16 hours with either 0.1% DMSO (negative control), 1 μg/mL Phytohemaglutinin (PHA) or 1µg/mL HIV overlapping peptide pool (synthesized by the MGH Peptide Core). The ELISpot assay was subsequently developed as per the ELISpot kit manufacturer’s instructions, and the results were reported as the number of IFN-γ spot forming cells (SFC) per 1 × 10^6^ PBMCs.

### Statistical analysis

In GRAd-COV2 toxicity studies run at CRL Evreux, body weight, rectal temperature, food consumption, CRP levels, hematology, blood biochemistry and coagulation data were analyzed using Provantis software. Shapiro-Wilk and Levene tests were used to assess respectively the normality and group homogeneity. When either test was significant, a log transformation was attempted, and the tests reapplied. The groups were compared using t-test if Shapiro-Wilk and Levene tests were not significant or the Wilcoxon test if either was significant. Organ weights were analyzed using PathData software. Kolmogorov and Bartlett tests were used to assess respectively the normality and group homogeneity. The groups were compared using t-test if Kolmogorov and Bartlett tests were not significant or the Wilcoxon test if either was significant.

For GRAdHIVNE1 toxicity study, all appropriate quantitative in-life data collected at AmplifyBio using the Provantis system were analyzed for test article effects by parametric or nonparametric analysis of variance (ANOVA). For all data, normality was determined by the Shapiro-Wilks test and homogeneity of variances was determined by Levene’s test. Data was log-transformed to meet parametric assumptions. For parametric data determined to be normally distributed and homogeneous among groups, an ANOVA F-test was used to determine whether there are differences among the group means. If the ANOVA F-test is significant, then tests for differences between the control and each of the comparison groups was conducted using Dunnett’s test. For nonparametric data that are not normally-distributed and/or nonhomogeneous, a Kruskal-Wallis test was used to determine whether there are differences among the group means. If the Kruskal-Wallis test is significant, then tests for differences between the control and each of the comparison groups was conducted using Wilcoxon tests and Bonferroni-Holm.

All statistical tests were performed at the 0.05 level of significance (p < 0.05).

The data obtained in the biodistribution studies were mostly descriptive and not analyzed statistically.

## Supporting information

Paalangara R Supplementary

## Data Availability

the dataset generated and analyzed for the current manuscript were extracted from the reports of proprietary, in some cases Good Laboratory Practice (GLP), Non-clinical Safety Studies. All relevant data are included in the article and in supplementary information file.

## Acknowledgments

The authors wish to thank the ReiThera PD, GMP and QC staff for the manufacturing and quality control of all vaccine preparations administered in the reported studies. The authors are grateful to Susan Barnett and Thandi M Onami, senior Program officers at the Gates Foundation, for their invaluable support. This research was funded by Regione Lazio and Italian Ministry of research (with no specific grant associated) for the GRAd-COV2 studies; and by the Gates Foundation, grants number INV-059646 (ReiThera), INV-008696 & INV-064567 (Ragon Institute) and INV-007375 & INV-065025 (IAVI) for the GRAdHIVNE1 studies. The conclusions and opinions expressed in this work are those of the author(s) alone and shall not be attributed to the Foundation. Under the grant conditions of the Foundation, a Creative Commons Attribution 4.0 License has already been assigned to the Author Accepted Manuscript version that might arise from this submission. Please note works submitted as a preprint have not undergone a peer review process.

## Author Contributions

conceptualization, R.P., J.A., A.F., S.Ca.; methodology, R.P., S.G., A.M., C.A., W.B., A.Ro., P.A., S.Ca.; formal analysis, S.G., A.M., C.A., W.B., A.Ro., M.A.G., A.A., S.B., G.D.G, P.A., S.Ca.; investigation, S.G., A.M., C.A., W.B., A.Ro., M.A.G., A.A., S.B.; resources, A.Ra., M.G., A.DR., A.N., G.M., F.G., D.S., F.N.; writing—original draft preparation, R.P., S.Ca.; writing—review and editing, S.G., A.M., C.A., W.B., A.Ro., M.A.G., B.C., G.D.G., J.A.; visualization, S.Ca.; supervision, R.P., A.T.M., P.A., A.F., S.Co., S.Ca.; project administration, M.S., B.C.; funding acquisition, A.F., S.Co.; All authors have read and agreed to the published version of the manuscript.

## Competing Interests

R.P., D.S. B.C. L.M. and V.M. are employees of IAVI; S.G., A.M., C.A., W.B., A.Ro. and P.A. are employees of Charles River Laboratories, a Contract Research Organization contracted by ReiThera and IAVI in the context of these studies; S.B., A.R., M.G., A.DR., A.N., G.M., F.G., F.N., M.S., A.F., S.Co and S.Ca. are or were at the time of the study, employees of the ReiThera SrL; A.F. and S.Co. are also founders and shareholders of Keires AG; A.R. and S.Co. are named inventors of the patent application no. 20183515.4 titled “Gorilla Adenovirus Nucleic Acid- and Amino Acid-Sequences, Vectors Containing Same, and Uses Thereof”; G.D.G. is a named inventor of patent application US20220323570A1 titled “Highly Networked Immunogen Composition”; G.D.G. receives research funding from Merck and Moderna; J.A. is paid consultant to IAVI; The other authors declare no conflict of interest. The funders had no role in the design of the study; in the collection, analyses, or interpretation of data; in the writing of the manuscript; or in the decision to publish the results.

