## Supplementary material for "Non-clinical safety of GRAd vector-based COVID-19 and HIV vaccines supports a platform regulatory approach": Paalangara R Supplementary

### Supplementary Information

Paalangara R. et al

**Supplementary table 1: GRAd toxicity and local tolerance studies in New Zealand White Rabbits**

| | **GRAd-COV2<br>Single Dose** | **GRAd-COV2<br>Repeated Dose** | **GRAdHIVNE1<br>Repeated Dose** |
|---|---|---|---|
| **age (weeks)<br>weight mean (sex, kg)** | 15-16w<br>M 3.1/F 3.3 | 13-18w<br>M 3.1/F 2.8 | 11w<br>M 2.2/F 2.1 |
| **N injections<br>interval** | N=1 (d1) | N=2<br>2 weeks (d1; d15) | N=2<br>3 weeks (d1; d22) |
| **vaccine dose** | 1 x $10^{11}$ vp (500 µL) | | |
| **follow up** | 1 week | 4 weeks post 2nd injection | |
| **group numerosity** | 5M+5F receiving vaccine or saline/euthanasia time point | | |
| **euthanasia time points** | d8 (7d post admin) | d18 (3d post 2nd)<br>d43 (4w post 2nd) | d25 (3d post 2nd)<br>d49F/d50M (4w post 2nd) |
| **vaccine take demonstration** | none | Spike Antibodies<br>(serum, ELISA) | T cells to HIV epitopes<br>(PBMC, IFNγ ELISpot) |
| **survival check** | Once in pretreatment, then twice daily | | |
| **clinical observation** | cageside: daily<br>detailed: once pre, d1@4h, d4, d7 | cageside: daily<br>detailed: once pre, d1@4h, then weekly | cageside: daily (on d1 and d22 @0.5h,3h, 8h)<br>detailed: d1 and d22@1h-6h,24h, then weekly |
| **body temperature** | pre (admin time/4h),<br>@admin(s), 4h, 24h<br>If >40°C daily until resolution | | 5x pre, d1-d22 @admin, 6h, 24h, 48h, 72h<br>If >40°C daily until resolution (2 occasions) |
| **body weight** | Thrice pre, admin days (d1) then daily until EOS | Thrice pre, admin days (d1 and d15), daily for next 3 days, then weekly | Once pre, d1, 2, 3, 8, 11, 15, 18, 21, 23, 24, then twice weekly. |
| **food consumption** | Once pre, d1 then daily | Once pre, d1, then twice weekly | d1, then daily |
| **local tolerance** | Draize score: @admin(s), 4h, 24h, 48h,72h | | Draize score: @admin, 6h, 24h, 48h,72h, EOS |
| **ophthalmology** | none | once pre, EOS | |
| **clinical pathology (hematology, biochemistry, coagulation, CRP)** | pre, d1(pre), d2, d4, d8 (EOS) | pre, @admin(pre), d2, d4, d8 after each admin, d43 (EOS) | pre, d2 (CRP only), d4, d25/d49 (EOS) |
| **anatomic pathology** | necropsy<br>organ weight,<br>preservation (limited list)<br>histopathology of inj. site and draining LN | necropsy<br>organ weight (selected list), preservation, histopathology (full WHO list) | |

*Supplementary table 1 abbreviations: h=hour; d=day; w=week; pre=pre-dose; admin=administration; EOS=end of study*

**Supplementary table 2: GRAd-COV2 Biodistribution Results (Genome Copies/µg of DNA)**

| Organ | Sex | Day 2 | | | Day 8 | | | Day 30 | | | Day 49 | | |
|---|---|---|---|---|---|---|---|---|---|---|---|---|---|
| | | No. of samples (/5) | | | No. of samples (/5) | | | No. of samples (/5) | | | No. of samples (/5) | | |
| | | BLD[a] | BLQ[b] | Quantified[c] | BLD[a] | BLQ[b] | Quantified[c] | BLD[a] | BLQ[b] | Quantified[c] | BLD[a] | BLQ[b] | Quantified[c] |
| Blood | F | 0 | 4 | 1<br>3.61E+02 | 5 | 0 | 0 | | | | | | |
| | M | 2 | 3 | 0 | 5 | 0 | 0 | | | | | | |
| Right Iliac LN | F | 0 | 0 | 5<br>Mean : 1.32E+04<br>(1.85E+03 - 3.99E+04) | 0 | 0 | 5<br>Mean : 1.23E+02<br>(8.20E+01 - 1.89E+02) | 0 | 3 | 2<br>Mean : 5.65E+01<br>(5.56E+01 - 5.74E+01) | 0 | 1 | 4<br>Mean : 1.24E+03<br>(7.96E+02 - 1.65E+03) |
| | M | 1 | 0 | 4<br>Mean : 1.55E+04<br>(1.99E+03 - 2.63E+04) | 0 | 0 | 5<br>Mean : 2.24E+03<br>(9.71E+01 - 5.45E+03) | 5 | 0 | 0 | 1 | 0 | 4<br>Mean : 1.16E+03<br>(8.98E+02 - 1.47E+03) |
| Right Inguinal LN | F | 3 | 0 | 2<br>Mean : 1.16E+03<br>(2.86E+02 - 2.02E+03) | 3 | 2 | 0 | 5 | 0 | 0 | 1 | 2 | 2<br>Mean : 8.26E+02<br>(6.60E+01 - 1.59E+03) |
| | M | 1 | 1 | 3<br>Mean : 2.71E+03<br>(4.98E+02 – 6.84E+03) | 1 | 1 | 3<br>Mean : 3.33E+02<br>(1.21E+02 - 7.27E+02) | 4 | 1 | 0 | 2 | 1 | 2<br>Mean : 1.53E+02<br>(1.00E+02 - 2.05E+02) |
| Quadriceps muscle | F | 0 | 0 | 5<br>Mean : 9.76E+04<br>(7.34E+02 - 3.84E+05) | 0 | 1 | 4<br>Mean : 6.69E+02<br>(9.38E+01 - 1.72E+03) | 0 | 0 | 5<br>Mean : 2.01E+02<br>(6.71E+01 - 4.63E+02) | 1 | 0 | 4<br>Mean : 5.82E+02<br>(8.93E+01 - 1.82E+03) |
| | M | 0 | 0 | 5<br>Mean : 1.15E+04<br>(7.53E+01 - 3.76E+04) | 0 | 0 | 5<br>Mean : 5.83E+03<br>(9.51E+01 - 1.13+04) | 2 | 1 | 2<br>Mean : 7.84E+01<br>(6.64E+01 - 9.05E+01) | 1 | 0 | 4<br>Mean : 6.24E+02<br>(9.77E+01 - 2.10E+03) |

[a] Below limit of detection (i.e., < 6.25 genome copies/well).

[b] Below limit of quantification (i.e., < 50 genome copies/well).

[c] Number of quantified samples and mean of test item DNA quantity (genome copies/µg of DNA) when applicable (i.e., unless only one sample returned a quantified value). In brackets: Min and Max values.

In grey = Analysis not performed as no test item genomic DNA was quantified at the two previous time points. LN = Lymph Nodes. No = Number

#### Supplementary table 3: GRAdHIVNE1 Biodistribution Results (Genome Copies/µg of DNA)

| Organ | Sex | Day 2 | | | Day 8 | | | Day 29 | | | Day 49 | | |
|---|---|---|---|---|---|---|---|---|---|---|---|---|---|
| | | No. of Samples (/5) | | | No. of Samples (/5) | | | No. of Samples (/5) | | | No. of Samples (/5) | | |
| | | BLD[a] | BLQ[b] | Quantified[c] | BLD[a] | BLQ[b] | Quantified[c] | BLD[a] | BLQ[b] | Quantified[c] | BLD[a] | BLQ[b] | Quantified[c] |
| Blood | F | 0 | 1 | 4<br>Mean: 1.99E+03<br>(3.03E+02-3.62E+03) | 5 | 0 | 0 | 4 | 1 | 0 | | | |
| | M | 0 | 2 | 3<br>Mean: 1.56E+02<br>(8.03E+01-3.03E+02) | 5 | 0 | 0 | 5 | 0 | 0 | | | |
| Iliac LN | F | 0 | 1 | 4<br>Mean: 9.78E+04<br>(6.21E+04-1.17E+05) | 0 | 0 | 5<br>Mean: 5.87E+03<br>(4.18E+03-7.15E+03) | 0 | 0 | 5<br>Mean: 5.09E+03<br>(3.75E+03-7.01E+03) | 1 | 0 | 4<br>Mean: 4.67E+03 (2.96E+03-6.35E+03) |
| | M | 0 | 1 | 4<br>Mean: 6.68E+04,<br>(5.33E+04-7.88E+04) | 0 | 0 | 5<br>Mean: 4.96E+03<br>(3.69E+03-7.33E+03) | 0 | 1 | 4<br>Mean: 2.88E+03<br>(1.90E+03-4.33E+03) | 1 | 1 | 3<br>Mean: 2.70E+03 (1.52E+03-3.47E+03) |
| Inguinal LN | F | 1 | 1 | 3<br>Mean: 1.53E+04<br>(1.64E+02-4.55E+04) | 0 | 1 | 4<br>Mean: 9.74E+02<br>(7.24E+01-1.91E+03) | 0 | 2 | 3<br>Mean:1.32E+02<br>(5.99E+01-2.67E+02) | 1 | 3 | 1<br>4.56E+03 |
| | M | 1 | 2 | 2<br>Mean:1.88E+04<br>(7.68E+01-3.74E+04) | 0 | 3 | 2<br>Mean:1.71E+03<br>(7.37E+01-3.34E+03) | 2 | 3 | 0 | 1 | 1 | 3<br>Mean: 1.32E+03 (1.01E+02-3.21E+03) |
| Quadriceps muscle | F | 0 | 0 | 5<br>Mean: 5.10E+06<br>(5.86E+05-1.43E+07) | 0 | 0 | 5<br>Mean: 5.00E+04<br>(9.34E+03-7.90E+04) | 0 | 0 | 5<br>Mean: 1.04E+04<br>(3.52E+02-1.42E+04) | 0 | 0 | 5<br>Mean: 7.27E+03 (2.08E+02-1.78E+04) |
| | M | 0 | 0 | 5<br>Mean: 2.08E+06<br>(5.04E+05-7.04E+06) | 0 | 0 | 5<br>Mean: 2.35E+04<br>(4.90E+02-4.60E+04) | 0 | 0 | 5<br>Mean: 4.44E+03<br>(3.04E+02-7.53E+03) | 0 | 0 | 5<br>Mean: 4.28E+03<br>(3.85E+02-1.02E+04) |
| Spleen | F | 0 | 2 | 3<br>Mean:1.96E+02<br>(5.04E+01-3.19E+02) | 0 | 5 | 0 | 0 | 2 | 3<br>Mean:9.79E+01<br>(6.72E+01-1.29E+02) | 0 | 3 | 2<br>Mean:1.26E+02 (1.22E+02-1.31E+02) |
| | M | 2 | 3 | 0 | 1 | 4 | 0 | 4 | 1 | 0 | 1 | 4 | 0 |

[a] Below limit of detection (i.e., < 6.25 genome copies/well).

[b] Below limit of quantification (i.e., < 50 genome copies/well).

[c] Number of quantified samples and mean of test item DNA quantity (genome copies/µg of DNA) when applicable (i.e., unless only one sample returned a quantified value). In brackets: Min and Max values.

In grey = Analysis not performed as no test item genomic DNA was quantified at the two previous time points. LN = Lymph Nodes. No = Number

#### Supplementary table 4: GRAd-COV2 RD toxicity study body temperature (°C)

| Sex | Male | | Female | |
|---|---|---|---|---|
| Group | 1 | 2 | 1 | 2 |
| Treatment | Control (Saline) | Vaccine (GRAd-COV2) | Control (Saline) | Vaccine (GRAd-COV2) |
| **1st administration: Day 1** | | | | |
| . Day 1 (pre-dose) | 39.0 | 38.9 | 39.1 | 39.1 |
| . Day 1 (4 hours) | 38.6 | 38.9 | 39.1 | 39.1 |
| . Day 2 (24 hours) | 38.5 | 39.0** | 38.6 | **39.5**** |
| . Day 3 (48 hours) | - | - | - | 38.8 (n = 3) |
| **2nd administration: Day 15** | | | | |
| . Day 15 (pre-dose) | 39.3 | 39.3 | 39.1 | 38.9 |
| . Day 15 (4 hours) | 38.7 | 38.7 | 38.8 | 39.1 |
| . Day 16 (24 hours) | 38.7 | 38.9 | 38.8 | 38.7 |

Statistically significant differences from controls: ** ($p<0.01$).

**Bold values**: considered as test item-related. -: not applicable; n: number of animals.

**Supplementary table 5: GRAd-COV2 RD toxicity study summary of Hematology Changes Compared to Control (Group 1) Values**

| Parameter/ Study Day | Males | | Females | |
|---|---|---|---|---|
| | Group 1 | Group 2 | Group 1 | Group 2 |
| | 0 vp/animal | 1 x $10^{11}$ vp/animal | 0 vp/animal | 1 x $10^{11}$ vp/animal |
| **White Blood cells (G/L)** | | | | |
| Pretest | 5.36 | 1.00x | 6.59 | 1.04x |
| Day 16 | 4.85 | **1.27x*** | 5.44 | **1.49x**** |
| Day 18 | 4.91 | **1.22x** | 7.36 | **1.11x** |
| **Neutrophils (G/L)** | | | | |
| Pretest | 1.13 | 1.09x | 1.22 | 1.18x |
| Day 2 | 0.83 | **1.64*** | 1.28 | **1.93x** |
| Day 16 | 0.78 | **2.20x**** | 0.95 | **2.70x***** |
| **Lymphocytes (G/L)** | | | | |
| Pretest | 3.62 | 0.93x | 4.55 | 1.02x |
| Day 2 | 3.55 | **0.83x*** | 3.84 | **0.68x*** |
| **Monocytes (G/L)** | | | | |
| Pretest | 0.11 | 1.27x | 0.15 | 0.73x |
| Day 4 | 0.11 | 1.55x | 0.19 | **2.16x** |
| Day 18 | 0.08 | 2.13x | 0.25 | **1.68x** |
| **Eosinophils (G/L)** | | | | |
| Pretest | 0.09 | 1.44x | 0.13 | 0.77x |
| Day 2 | 0.12 | **0.50x*** | 0.08 | **0.38x**** |
| **Basophils (G/L)** | | | | |
| Pretest | 0.39 | 1.10x | 0.51 | 1.06x |
| Day 2 | 0.41 | **0.80x** | 0.46 | **0.76x** |
| **Large unstained cells (G/L)** | | | | |
| Pretest | 0.03 | 0.67x | 0.03 | 0.67x |
| Day 16 | 0.02 | 1.00x | 0.03 | **1.67x** |
| Day 18 | 0.02 | **2.50x** | 0.03 | **2.33x** |
| **Platelets (G/L)** | | | | |
| Pretest | 386 | 0.83x | 397 | 1.04x |
| Day 2 | 410 | **0.79x** | 406 | **0.76x** |

Vp = viral particle, Statistically significant from controls: * (p<0.05); ** (p<0.01); *** (p<0.001).
**Bold values**: considered as test item-related;
The fold change (x) of test item-related findings relative to the control values are listed. For comparison, the control (Group 1) are listed.

**Supplementary table 6: GRAd-COV2 RD toxicity study summary of Coagulation Changes Compared to Control (Group 1) Values**

| Parameter/ Study Day | Males | | Females | |
|---|---|---|---|---|
| | Group 1 | Group 2 | Group 1 | Group 2 |
| | 0 vp/animal | 1 x $10^{11}$ vp/animal | 0 vp/animal | 1 x $10^{11}$ vp/animal |
| **Fibrinogen (g/L)** | | | | |
| Pretest | 2.85 | 0.98x | 2.02 | 0.91x |
| Day 2 | 2.96 | **1.43x**** | 1.99 | **1.97x***** |
| Day 4 | 2.81 | **1.35x**** | 1.87 | **1.85x***** |
| Day 16 | 2.58 | **1.51x***** | 1.93 | **2.02***** |
| Day 18 | 2.65 | **1.23x**** | 1.91 | **1.48*** |

Vp = viral particle, Statistically significant from controls: * (p<0.05); ** (p<0.01); *** (p<0.001).
**Bold values**: considered as test item-related;
The fold change (x) of test item-related findings relative to the control values are listed. For comparison, the control (Group 1) are listed.

**Supplementary table 7: GRAd-COV2 RD toxicity study summary of Blood Biochemistry Changes Compared to Control (Group 1) Values**

| Parameter/ Study Day | Males | | Females | |
|---|---|---|---|---|
| | Group 1 | Group 2 | Group 1 | Group 2 |
| | 0 vp/animal | 1 x $10^{11}$ vp/animal | 0 vp/animal | 1 x $10^{11}$ vp/animal |
| **Albumin (g/L)** | | | | |
| Pretest | 45 | 0.97x | 41 | 1.05x |
| Day 16 | 44 | 1.00x | 40 | 1.05x |
| Day 22 | 44 | 1.02x | 41 | 1.02x |
| **A/G** | | | | |
| Pretest | 3.41 | 0.92 | 3.58 | 1.06 |
| Day 16 | 3.44 | **0.87x*** | 3.60 | **0.92x*** |
| Day 22 | 3.50 | **0.87x**** | 4.00 | 0.90x |

Vp = viral particle, Statistically significant from controls: * (p<0.05); ** (p<0.01); *** (p<0.001).
**Bold values**: considered as test item-related;
The fold change (x) of test item-related findings relative to the control values are listed. For comparison, the control (Group 1) are listed.

**Supplementary table 8: GRAd-COV2 RD toxicity study summary of C-Reactive Protein (CRP) Levels Changes Compared to Control (Group 1) Values**

| Parameter/ Study Day | Males | | Females | |
|---|---|---|---|---|
| | Group 1 | Group 2 | Group 1 | Group 2 |
| | 0 vp/animal | 1 x $10^{11}$ vp/animal | 0 vp/animal | 1 x $10^{11}$ vp/animal |
| **CRP (µg/mL)** | | | | |
| Pretest | 2.24 | 2.41x | 5.93 | 1.12x |
| Day 2 | 4.10 | **23.31x***** | 7.44 | **12.01***** |
| Day 16 | 1.86 | **37.19x*** | 9.79 | **10.50*** |

Vp = viral particle, Statistically significant from controls: * (p<0.05); ** (p<0.01); *** (p<0.001).
**Bold values**: considered as test item-related;
The fold change (x) of test item-related findings relative to the control values are listed. For comparison, the control (Group 1) are listed.

**Supplementary table 9: GRAd-COV2 RD toxicity study summary of Treatment-related Findings in Mean Organ Weights Compared to Control (Group 1) Values at Early Euthanasia**

| | **Male**<br>**Dosage (vp/animal/adm.)** | **Female**<br>**Dosage (vp/animal/adm.)** |
|---|---|---|
| | **$1 \times 10^{11}$** | **$1 \times 10^{11}$** |
| **Spleen** | | |
| .absolute | +42 | +48* |
| .relative to body weight | +52 | +48* |
| .relative to brain weight | +53 | +55* |
| **Right iliac lymph node** | | |
| .absolute | +165 | +194* |
| .relative to body weight | +178 | +194* |
| .relative to brain weight | +184 | +210* |
| **Right popliteal lymph node** | | |
| .absolute | +65** | +67 |
| .relative to body weight | +79** | +63 |
| .relative to brain weight | +77** | +77 |
| **Right inguinal lymph node** | | |
| .absolute | +63 | -13 |
| .relative to body weight | +70 | -12 |
| .relative to brain weight | +68 | -8 |
| **Adrenal glands** | | |
| .absolute | +23 | +14 |
| .relative to body weight | +31* | +15 |
| .relative to brain weight | +32* | +21* |
| **Thymus** | | |
| .absolute | -25 | -14 |
| .relative to body | -19 | -16 |
| .relative to brain | -21 | -9 |

Statistically significant from controls: *: p<0.05, **: p<0.01.

The significance concerned the organ weights values and not the percentages.

**Supplementary table 10: GRAd-COV2 RD toxicity study summary of Treatment-related Findings in Mean Organ Weights Compared to Control (Group 1) Values at Late Euthanasia**

| | Male Dosage (vp/animal/adm.) | Female Dosage (vp/animal/adm.) |
|---|---|---|
| | $1 \times 10^{11}$ | $1 \times 10^{11}$ |
| **Right iliac lymph node** | | |
| .absolute | +118* | -30 |
| .relative to body weight | +97* | -25 |
| .relative to brain weight | +125* | -29 |
| **Right popliteal lymph node** | | |
| .absolute | +26* | +28 |
| .relative to body weight | +15 | +33 |
| .relative to brain weight | +28* | +26 |
| **Left iliac lymph node** | | |
| .absolute | +58 | -8 |
| .relative to body weight | +46 | +6 |
| .relative to brain weight | +62 | -8 |

Statistically significant from controls: *: $p<0.05$

The significance concerned the organ weights values and not the percentages.

**Supplementary table 11: GRAd-COV2 RD toxicity study summary of Treatment-related Macroscopic Findings Compared to Control (Group 1) Values at Early Euthanasia**

| Finding | Male<br>Dosage (vp/animal/adm.) | | Female<br>Dosage (vp/animal/adm.) | |
|---|---|---|---|---|
| | **0** | **1 x $10^{11}$** | **0** | **1 x $10^{11}$** |
| **Right iliac lymph node** (number examined) | 5 | 5 | 5 | 5 |
| Enlarged | - | 2 | - | 4 |
| **Left iliac lymph node** (number examined) | 5 | 5 | 5 | 5 |
| Enlarged | - | - | - | 1 |

-: finding not present.

**Supplementary table 12: GRAd-COV2 RD toxicity study summary of Treatment-related Microscopic Findings Compared to Control (Group 1) Values at Early Euthanasia**

| Finding | Male Dosage (vp/animal/adm.) | | Female Dosage (vp/animal/adm.) | |
|---|---|---|---|---|
| | 0 | 1 x $10^{11}$ | 0 | 1 x $10^{11}$ |
| **Spleen** (*number examined*) | *5* | *5* | *5* | *5* |
| Cellularity: increased; lymphoid | | | | |
| Minimal (grade 1) | - | 3 | - | 1 |
| Slight (grade 2) | - | 2 | - | 2 |
| Moderate (grade 3) | - | - | - | 2 |
| **Right iliac lymph node** (*number examined*) | *5* | *4* | *5* | *5* |
| Cellularity: increased; lymphoid | | | | |
| Slight (grade 2) | - | 2 | - | - |
| Moderate (grade 3) | - | 2 | - | 2 |
| Marked (grade 4) | - | - | - | 3 |
| Intrasinusoidal erythrocytes | | | | |
| Minimal (grade 1) | - | 3 | 1 | 2 |
| **Right inguinal lymph node** (*number examined*) | *5* | *4* | *4* | *5* |
| Cellularity: increased; lymphoid | | | | |
| Minimal (grade 1) | - | 1 | - | 1 |
| Intrasinusoidal erythrocytes | | | | |
| Minimal (grade 1) | - | - | 1 | - |
| **Right popliteal lymph node** (*number examined*) | *5* | *5* | *5* | *5* |
| Cellularity: increased; lymphoid | | | | |
| Minimal (grade 1) | - | 2 | - | - |
| Slight (grade 2) | - | 1 | - | 4 |
| Erythrophagocytosis | | | | |
| Minimal (grade 1) | - | - | - | 1 |
| **Left iliac lymph node** (*number examined*) | *4* | *5* | *5* | *5* |
| Cellularity: increased; lymphoid | | | | |
| Minimal (grade 1) | - | 1 | - | - |
| Slight (grade 2) | - | 1 | - | - |
| Moderate (grade 3) | - | 2 | - | 1 |
| Intrasinusoidal erythrocytes | | | | |

| | | | | |
|---|---|---|---|---|
| Minimal (grade 1) | - | - | - | 1 |
| Slight (grade 2) | - | 1 | - | 1 |
| **Left inguinal lymph node** (*number examined*) | *5* | *4* | *4* | *4* |
| Cellularity: increased; lymphoid | | | | |
| Minimal (grade 1) | - | - | - | 1 |
| **Left popliteal lymph node** (*number examined*) | *5* | *5* | *5* | *5* |
| Cellularity: increased; lymphoid | | | | |
| Minimal (grade 1) | - | - | - | 2 |
| **Injection site 2** (*number examined*) | *5* | *5* | *5* | *5* |
| Infiltrate; mononuclear inflammatory cell infiltrate | | | | |
| Minimal (grade 1) | - | 3 | - | 2 |
| Infiltrate; mixed inflammatory cell infiltrate | | | | |
| Minimal (grade 1) | - | 1 | - | - |
| Slight (grade 2) | - | - | - | 1 |
| **Sciatic nerve** (*number examined*) | *5* | *5* | *4* | *5* |
| Infiltrate; mononuclear inflammatory cell infiltrate | | | | |
| Minimal (grade 1) | - | 1 | - | - |
| Slight (grade 2) | - | 1 | - | 4 |
| Infiltrate; mixed inflammatory cell infiltrate | | | | |
| Minimal (grade 1) | - | 1 | - | - |
| Slight (grade 2) | - | 1 | - | - |
| **Femur** (*number examined*) | *5* | *5* | *5* | *5* |
| Infiltrate; mixed inflammatory cell infiltrate | | | | |
| Minimal (grade 1) | - | - | - | 1 |

-: finding not present.

**Supplementary table 13: GRAd-COV2 RD toxicity study summary of Treatment-related Microscopic Findings Compared to Control (Group 1) Values at Late Euthanasia**

| Finding | Male Dosage (vp/animal/adm.) | | Female Dosage (vp/animal/adm.) | |
|---|---|---|---|---|
| | 0 | 1 x $10^{11}$ | 0 | 1 x $10^{11}$ |
| **Spleen** (*number examined*) | *5* | *5* | *5* | *5* |
| Cellularity: increased; lymphoid | | | | |
| Minimal (grade 1) | - | 4 | - | 4 |
| **Right iliac lymph node** (*number examined*) | *5* | *5* | *5* | *5* |
| Cellularity: increased; lymphoid | | | | |
| Slight (grade 2) | - | 2 | - | 4 |
| Moderate (grade 3) | - | 3 | - | - |
| **Right popliteal lymph node** (*number examined*) | *5* | *5* | *5* | *5* |
| Cellularity: increased; lymphoid | | | | |
| Minimal (grade 1) | - | 1 | - | 4 |
| Slight (grade 2) | - | 3 | - | - |
| **Left iliac lymph node** (*number examined*) | *5* | *4* | *5* | *5* |
| Cellularity: increased; lymphoid | | | | |
| Minimal (grade 1) | - | - | 1 | - |
| Slight (grade 2) | - | - | - | 1 |
| Moderate (grade 3) | - | 2 | - | 2 |
| **Left popliteal lymph node** (*number examined*) | *5* | *5* | *5* | *5* |
| Cellularity: increased; lymphoid | | | | |
| Minimal (grade 1) | - | - | - | 3 |

-: finding not present.

#### Supplementary table 14: GRAdHIVNE1 RD toxicity study body temperature (°C)

| **Sex: male** | study day | -7 → -1 | -6 | -5 | -4 | -3 | -2 | 1 (PD) | 1 (6h PE) | 2 (24h PE) | 3 (48h PE) | 4 (72h PE) | 22 (PD) | 22 (6h PE) | 23 (24h PE) | 24 (48h PE) | 25 (72h PE) |
|---|---|---|---|---|---|---|---|---|---|---|---|---|---|---|---|---|---|
| Group 1 | Mean | 38.2 | 38.3 | 38.0 | 38.2 | 38.2 | 38.3 | 38.3 | 38.5 | 38.3 | 38.5 | 38.3 | 38.6 | 38.9 | 39.0 | 39.0 | 38.3 |
| 0 | SD | 0.2 | 0.4 | 0.3 | 0.3 | 0.3 | 0.3 | 0.5 | 0.3 | 0.3 | 0.4 | 0.4 | 0.2 | 0.2 | 0.3 | 0.2 | 0.5 |
| vp/dose | N | . | 8 | 8 | 8 | 8 | 8 | 8 | 8 | 8 | 8 | 8 | 8 | 8 | 8 | 8 | 8 |
| Group 2 | Mean | 38.2 | 38.3 | 38.3 | 38.2 | 38.0 | 38.2 | 38.3 | 38.6 | 39.4 * | 38.9 | 38.0 | 38.5 | 39.0 | 38.6 | 38.7 | 38.3 |
| 1.0E+11 | SD | 0.4 | 0.7 | 0.4 | 0.4 | 0.4 | 0.6 | 0.3 | 0.4 | 0.5 | 0.6 | 0.5 | 0.3 | 0.4 | 0.5 | 0.4 | 0.6 |
| vp/dose | N | . | 9 | 10 | 10 | 10 | 10 | 10 | 10 | 10 | 10 | 10 | 10 | 10 | 10 | 10 | 10 |

| **Sex: Female** | study day | -7 → -1 | -7 | -6 | -5 | -4 | -3 | 1 (PD) | 1 (6h PE) | 2 (24h PE) | 3 (48h PE) | 4 (72h PE) | 22 (PD) | 22 (6h PE) | 23 (24h PE) | 24 48h PE) | 25 (72h PE) |
|---|---|---|---|---|---|---|---|---|---|---|---|---|---|---|---|---|---|
| Group 1 | Mean | 38.4 | 38.5 | 38.4 | 38.3 | 38.5 | 38.5 | 38.6 | 38.7 | 38.4 | 38.5 | 38.3 | 38.5 | 38.9 | 39.3 | 39.0 | 38.3 |
| 0 | SD | 0.2 | 0.5 | 0.3 | 0.3 | 0.3 | 0.2 | 0.4 | 0.2 | 0.3 | 0.3 | 0.4 | 0.4 | 0.4 | 0.2 | 0.2 | 0.6 |
| vp/dose | N | . | 8 | 8 | 8 | 8 | 8 | 8 | 8 | 8 | 8 | 8 | 8 | 8 | 8 | 8 | 8 |
| Group 2 | Mean | 38.5 | 38.5 | 38.6 | 38.6 | 38.5 | 38.5 | 38.6 | 38.9 | 40.3 * | 39.1 * | 38.8 | 38.7 | 39.2 | 39.1 | 39.0 | 38.1 |
| 1.0E+11 | SD | 0.4 | 0.6 | 0.5 | 0.5 | 0.4 | 0.4 | 0.3 | 0.4 | 0.5 | 0.5 | 0.5 | 0.3 | 0.4 | 0.2 | 0.2 | 0.5 |
| vp/dose | N | . | 10 | 10 | 10 | 10 | 10 | 10 | 10 | 10 | 10 | 10 | 10 | 10 | 10 | 10 | 10 |

Anova & Dunnett: * = $p < 0.05$

**Supplementary table 15: GRAdHIVNE1 RD toxicity study summary of Hematology Changes Compared to Control (Group 1) Values**

| Parameter/ Study Day | Males | | Female | |
|---|---|---|---|---|
| | Group 1 | Group 2 | Group 1 | Group 2 |
| | 0 vp/dose | 1.0E+11 vp/dose | 0 vp/dose | 1.0E+11 vp/dose |
| **Red Blood Cells (× $10^6$ cells/µL)** | | | | |
| Day 4 | 6.03 | **0.9×**[a] | 5.57 | **0.9×**[a] |
| Day 25 | 6.09 | **0.9×**[a] | 5.41 | 1.0× |
| Day 49/50 | 6.00 | 1.0× | 5.31 | 1.1× |
| **Hemoglobin (g/dL)** | | | | |
| Day 4 | 12.3 | **0.9×**[a] | 11.3 | **0.9×**[a] |
| Day 25 | 12.3 | **0.9×**[a] | 10.9 | 1.0× |
| Day 49/50 | 12.3 | 1.0× | 11.0 | 1.0× |
| **Hematocrit (%)** | | | | |
| Day 4 | 37.2 | **0.9×**[a] | 34.7 | **0.9×**[a] |
| Day 25 | 37.7 | **0.9×**[a] | 34.0 | 1.0× |
| Day 49/50 | 37.2 | 1.0× | 33.2 | 1.1× |
| **Monocytes (× $10^3$ cells/µL)** | | | | |
| Day 4 | 0.05 | **10.8×**[a] | 0.09 | **4.7×**[a] |
| Day 25 | 0.09 | **2.6×**[a] | 0.07 | **3.3×**[a] |
| Day 49/50 | 0.04 | 0.8× | 0.06 | 0.8× |
| **Monocytes (%)** | | | | |
| Day 4 | 0.65 | **11.7×**[a] | 1.50 | **4.6×**[a] |
| Day 25 | 1.43 | **2.1×**[a] | 1.58 | **2.7×**[a] |
| Day 49/50 | 0.98 | 0.7× | 1.58 | 0.9× |
| **Large Unstained Cells (× $10^3$ cells/µL)** | | | | |
| Day 4 | 0.01 | **3.0×**[a] | 0.01 | **2.0×**[a] |
| Day 25 | 0.01 | **2.0×**[a] | 0.00 | **-**[a] |
| Day 49/50 | 0.00 | - | 0.00 | - |
| **Large Unstained Cells (%)** | | | | |
| Day 4 | 0.1 | **4.0×**[a] | 0.2 | **2.0×**[a] |
| Day 25 | 0.1 | **3.0×**[a] | 0.1 | **2.0×**[a] |
| Day 49/50 | 0.1 | 1.0× | 0.1 | 1.0× |
| **Mean Platelet Volume (fL)** | | | | |
| Day 4 | 10.3 | **1.1×**[a] | 9.1 | **1.0×**[a] |
| Day 25 | 9.1 | 1.0× | 9.1 | 1.0× |
| Day 49/50 | 10.2 | 1.0× | 9.1 | 1.0× |

vp = viral particle, a = $p < 0.05$, - = no calculatable fold change

The fold change (×) of test article-related findings relative to the control values are listed. For comparison, the control (Group 1) values are listed.

**Supplementary table 16: GRAdHIVNE1 RD toxicity study summary of Serum Chemistry Changes Compared to Control (Group 1) Values**

| Parameter/ Study Day | Males | | Female | |
|---|---|---|---|---|
| | **Group 1** | **Group 2** | **Group 1** | **Group 2** |
| | **0 vp/dose** | **1.0E+11 vp/dose** | **0 vp/dose** | **1.0E+11 vp/dose** |
| **C-Reactive Protein (mg/L)** | | | | |
| Pre-dose | < | < | 30.5 | < |
| Day 2 | 18.0 | **4.1×[a]** | 21.2 | **4.7×[a]** |
| Day 4 | 8.7 | **7.3×[a]** | 18.6 | **3.4×[a]** |
| Day 25 | < | - | < | - |
| Day 49/50 | 7.6 | 1.0× | < | - |
| **Creatine Kinase (U/L)** | | | | |
| Day 4 | 1658 | > | > | > |
| Day 25 | 871 | **1.6×[a]** | 1172 | 1.1× |
| Day 49/50 | 1069 | 1.2× | > | - |
| **Rabbit Globulin (g/dL)** | | | | |
| Day 4 | 1.4 | **1.1×[a]** | 1.3 | **1.2×[a]** |
| Day 25 | 1.3 | **1.2×[a]** | 1.1 | **1.3×[a]** |
| Day 49/50 | 1.3 | 1.0× | 1.2 | 1.0× |
| **Hemolysis Index** | | | | |
| Day 4 | 7 | **0.6×[a]** | 4 | 1.5× |
| Day 25 | 14 | 0.7× | 13 | 0.5× |
| Day 49/50 | 11 | 1.1× | 12 | 0.9× |
| **Lipemia Index** | | | | |
| Day 4 | 7 | 1.0× | 5 | 1.2× |
| Day 25 | 2 | 1.5× | 1 | **4.0×[a]** |
| Day 49/50 | 1 | 2.0× | 1 | 2.0× |

vp = viral particle; a = $p < 0.05$; - = No calculated data available due to linearity exclusion.
< = Result below instrument linearity limit, too low to enumerate accurately. > = Result above instrument linearity limit, too high to enumerate accurately. The fold change (×) of test article-related findings relative to the control values are listed. For comparison, the control (Group 1) values are listed.

**Supplementary table 17: GRAdHIVNE1 RD toxicity study summary of Coagulation Changes Compared to Control (Group 1) Values**

| | Males | | Female | |
|---|---|---|---|---|
| | Group 1 | Group 2 | Group 1 | Group 2 |
| | 0 vp/dose | 1.0E+11 vp/dose | 0 vp/dose | 1.0E+11 vp/dose |
| **Prothrombin Time (seconds)** | | | | |
| Day 4 | 8.8 | 1.0× | 9.0 | **0.9×**[a] |
| Day 25 | 8.9 | **1.0×**[a] | 8.9 | **0.9×**[a] |
| Day 49/50 | 9.0 | 1.0× | 8.6 | 1.0× |
| **Activated Partial Thromboplastin Time (seconds)** | | | | |
| Day 4 | 14.1 | 1.0× | 13.8 | **0.9×**[a] |
| Day 25 | 14.7 | 0.9× | 15.6 | **0.9×**[a] |
| Day 49/50 | 15.1 | 1.1× | 16.5 | 1.0× |
| **Fibrinogen (mg/dL)** | | | | |
| Day 4 | 355 | **1.7×**[a] | 301 | **1.7×**[a] |
| Day 25 | 295 | **1.5×**[a] | 251 | **1.6×**[a] |
| Day 49/50 | 275 | 0.9× | 216 | 1.0× |

vp = viral particle, a = $p < 0.05$

The fold change (×) of test article-related findings relative to the control values are listed. For comparison, the control (Group 1) values are listed.

**Supplementary table 18: GRAdHIVNE1 RD toxicity study Summary of Treatment-Related Findings in the Injection Sites and Skeletal Muscle – Main Cohort (Day 25)**

| | Males | | Females | |
|---|---|---|---|---|
| **Group** | **1** | **2** | **1** | **2** |
| **Dose (vp/dose)** | **0** | **1.0E+11** | **0** | **1.0E+11** |
| **No. of animals examined** | **4** | **5** | **4** | **5** |
| **Injection site 1 (no. examined)** | 4 | 5 | 4 | 5 |
| Inflammation, skeletal muscle, mononuclear cell | 0 | 2 (1.5) | 0 | 1 (2.0) |
| Degeneration, skeletal muscle | 0 | 1 (1.0) | 0 | 0 |
| **Injection site 2 (no. examined)** | 4 | 5 | 4 | 5 |
| Inflammation, skeletal muscle, acute | 0 | 0 | 0 | 1 (2.0) |
| Hemorrhage, skeletal muscle | 0 | 0 | 0 | 1 (2.0) |
| Necrosis, skeletal muscle | 0 | 2 (1.0) | 0 | 1 (2.0) |
| Regeneration, skeletal muscle | 0 | 0 | 0 | 1 (1.0) |
| Hemorrhage, dermis | 0 | 0 | 0 | 1 (2.0) |
| **Skeletal muscle** | 4 | 5 | 4 | 5 |
| Inflammation, mononuclear cell | 0 | 1 (1.0) | 0 | 1 (1.0) |
| Necrosis | 0 | 0 | 1 (2.0) | 1 (2.0) |
| Regeneration | 0 | 1 (1.0) | 0 | 0 |

Average severity (in parentheses) was calculated by adding all the grades of affected animals in the group and dividing by the number affected.

**Supplementary table 19: GRAdHIVNE1 RD toxicity study Summary of Treatment-Related Findings in the Injection Site 2 and Skeletal Muscle – Recovery Cohort (Day 49/50)**

| | Males | | Females | |
|---|---|---|---|---|
| **Group** | **1** | **2** | **1** | **2** |
| **Dose (vp/dose)** | 0 | 1.0E+11 | 0 | 1.0E+11 |
| **No. of animals examined** | 4 | 5 | 4 | 5 |
| **Injection site 2 (no. examined)** | 4 | 5 | 4 | 5 |
| Inflammation, skeletal muscle, mononuclear cell | 0 | 3 (1.3) | 0 | 0 |
| **Skeletal muscle** | 4 | 5 | 4 | 5 |
| Inflammation, mononuclear cell | 0 | 3 (1.0) | 0 | 1 (1.0) |

Average severity (in parentheses) was calculated by adding all the grades of affected animals in the group and dividing by the number affected.

**Supplementary table 20: Toxicology and Biodistribution studies of replication deficient adenoviral vectors by intramuscular route**

| Vector (Ad species) origin | Transgene pathogen/antigen | Ad genomic deletions | Type of study | Species and strain | Vaccine dose | Vaccine regimen | Reference |
|---|---|---|---|---|---|---|---|
| GRAd32 (C) gorilla | SARS-CoV-2/Spike glycoprotein | ΔE1-ΔE3-ΔE4 | SD Toxicity | Rabbit NZ | $1 \times 10^{11}$vp | d1 | this paper |
| | | | RD Toxicity | Rabbit NZ | $1 \times 10^{11}$ vp | d1-d15 | |
| | | | Biodistribution | Rat SD | $2 \times 10^{10}$ vp | d1 | |
| GRAd32 (C) gorilla | HIV/Polyepitope string | ΔE1-ΔE3 | RD Toxicity | Rabbit NZ | $1 \times 10^{11}$ vp | d1-d21 | this paper |
| | | | Biodistribution | Rat SD | $3.3 \times 10^{10}$ vp | d1 | |
| ChAd155 (C) chimpanzee | Rabies virus/ G glycoprotein | ΔE1-ΔE4 | Biodistribution | Rat SD | $2.3 \times 10^{10}$ vp | d1 | *Napolitano F, PLOS NTD 2020* |
| ChAd155 (C) chimpanzee | Respiratory Syncytial virus/F-N-M2-1 | ΔE1-ΔE4 | RD Toxicity | Rabbit NZ | $5 \times 10^{10}$ vp | d1-d15-d29 | *Stokes AH, IntJTox 2022* |
| | | | Biodistribution | Rat SD | $1 \times 10^{10}$ vp | d1 | |
| | | | Shedding | Rat SD | $1 \times 10^{10}$ vp | d1 | |
| ChAd3 (C) chimpanzee | Ebola/GP glycoprotein | ΔE1-ΔE4 | RD Toxicity | Rabbit NZ | $1.6 \times 10^{11}$ vp | d1-d22 | *Planty C, J.Appl.Tox 2020* |
| | | | Biodistribution | Rat SD | $1.6 \times 10^{10}$ vp | d1 | |
| ChAdOx1 (E) chimpanzee | SARS-CoV-2/Spike glycoprotein | ΔE1-ΔE3 | RD Toxicity | Mice CD-1 | $3.7 \times 10^{10}$ vp | d1 –d22-d43 | AZ COVID vax EMA/94907/2021 |
| | | | Biodistribution | Mice CD-1 | $7 \times 10^{9}$ vp | d1 | *Stebbings R, Vaccine 2022* |
| AdC68 (E) chimpanzee | SARS-CoV-2/Spike glycoprotein | ΔE1-ΔE3 | RD Toxicity | Rat SD/NHP Cyno | $2 \times 10^{11}$ vp /$4 \times 10^{11}$ vp | d1 - d15 (d19) | *Dai X, Immunotoxicology 2022* |
| | | | Biodistribution | Rat SD/NHP Cyno | $2 \times 10^{11}$ vp /$4 \times 10^{11}$ vp | d1 | |
| Ad26 (D) human | SARS-CoV-2/Spike glycoprotein | ΔE1-ΔE3 | RD Toxicity | Rabbit NZ | $1 \times 10^{11}$ vp | d1-d15-d29 | J&J COVID vax EMA/158424/2021 |
| Ad5 (C) human | SARS-CoV-2/Spike glycoprotein | ΔE1-ΔE3 | Biodistribution | Mice BALB/c | $1 \times 10^{8}$ IFU | d1 | *Dong Lee H, J.Microbiol.Biotechnol 2023* |
| several Ad5 (C) human | HIV/gag-pol, env Ebola or Marburg/GP glycoprotein | ΔE1-ΔE3 or ΔE1-ΔE3-ΔE4 | RD Toxicity | Rabbit NZ | 1-2 $\times 10^{11}$ vp/PU | d1 –d22 (d43) | *Sheets RL, J.Immunotox 2008* |
| Ad35 (B) human | HIV/env | ΔE1 | Biodistribution | Rabbit NZ | 0.5-1 $\times 10^{11}$ vp/PU | d1 | |
| Ad5/35 chimera (C/B) human | HIV/gag | ΔE1-ΔE3 | Biodistribution | Mice BALB/c | $2 \times 10^{9}$ vp | d1 | *Shimada M, gene Therapy 2022* |

*Supplementary Table 20 Abbreviations: SD Tox= single dose; RD Tox=repeated dose; NZ=New Zealand; SD (referred to Rats)= Sprague Dawley; NHP=non-human primate; Cyno= Cynomolgus macaque*
